# Mammalian PIWI-piRNA-target complexes reveal features for broad and efficient target-silencing

**DOI:** 10.1101/2023.06.23.546240

**Authors:** Zhiqing Li, Zhenzhen Li, Lili Li, Lin Zeng, Junchao Xue, Huilin Niu, Jing Zhong, Qilu Yu, Dengfeng Li, Shikui Tu, ZZ Zhao Zhang, Chun-Qing Song, Jianping Wu, En-Zhi Shen

## Abstract

The PIWI-interacting RNA (piRNA) pathway is an adaptive defense system wherein piRNAs guide PIWI-family Argonaute proteins to recognize and silence ever-evolving selfish genetic elements and ensure genome integrity. Driven by this intensive host-pathogen arms race, piRNA pathway and its targeted transposons have co-evolved rapidly in a species-specific manner, but how the piRNA pathway specifically adapts to target silencing in mammals remains elusive. Here, we show that mouse MILI and human HILI piRISCs bind and cleave targets more efficiently than their invertebrate counterpart from sponge *Ephydatia fluviatilis*. The inherent functional differences comport with structural features identified by cryo-EM studies of piRISCs. In the absence of target, MILI and HILI piRISCs adopt a wider nucleic-acid-binding channel and display an extended prearranged piRNA seed as compared to *Ef*Piwi piRISC, consistent with their ability to capture targets more efficiently than *Ef*Piwi piRISC. In the presence of target, the seed gate—which enforces seed-target fidelity in microRNA RISC—adopts a relaxed state in mammalian piRISC, revealing how MILI and HILI tolerate seed-target mismatches to broaden the target spectrum. A vertebrate-specific lysine distorts the piRNA seed, shifting the trajectory of the piRNA-target duplex out of the central cleft and toward the PAZ lobe. Functional analyses reveal that this lysine promotes target binding and cleavage. Our study therefore provides a molecular basis for the piRNA targeting mechanism in mouse and human, and suggests that mammalian piRNA machinery can achieve broad target silencing using a limited supply of piRNA species.

## Main Text

Transposons, referred to as genomic ‘pathogens’, are able to mobilize and replicate themselves within a genome^1^. They evolve rapidly and act as a powerful force for genomic evolution by fueling the development of new proteins and activities^2–8^. Despite these beneficial effects, uncontrolled transposon mobilization causes genetic instability, sterility, and disease. In animals, PIWI proteins and their small RNA guides (piRNAs) maintain genome integrity and germ cell development to ensure fertility and the faithful transmission of genetic information across generations ^9–13^. Whereas transposons—the primary targets of piRNA-directed silencing—occupy a large fraction of the genome in most mammals, the source loci of piRNAs (called piRNA clusters) occupy a minute fraction^11, 14–20^. For example, retrotransposons comprise at least 45% of the human genome^15, 17^, but piRNA clusters occupy only ∼0.37% of the human genome^20^ (**Extended Data Fig. 1**). By contrast, the transposon to piRNA cluster content is markedly lower in the *Drosophila melanogaster* genome (14% to transposon, 3.2% to piRNA clusters)^15, 17, 20^. As a rapidly evolving pathway to cope with the ever-changing transposon dynamics within the genome, the piRNA silencing machinery likely has co-evolved with its host genomic parasites across different species. Therefore, mammalian PIWI proteins may possess distinct targeting features from invertebrates to meet their roles of guarding the mammalian germline genome.

Structural studies of PIWI proteins from invertebrates, including silkworm, fly, and sponge, reveal a bilobed architecture, in which N-PAZ and MID-PIWI lobes form the wall of the nucleic-acid-binding channel. These invertebrate PIWIs prearrange a short seed segment and have a lower binding affinity than miRNA targeting^21–23^. In addition, a small α-helix (∼1.5-turn) extends the ’seed gate’ structure and widens the channel near the piRNA seed region, which enforces seed-match targeting and facilitates subsequent propagation of piRNA-target base pairing to activate catalysis. Although they are expected to be similar, structures of vertebrate PIWI complexes— especially human PIWI complexes—have not been reported.

We have purified mouse and human Ago3-like PIWI proteins, MILI and HILI, and performed structure-function analyses to compare to invertebrate PIWI. We show that MILI and HILI piRNA-induced silencing complexes (piRISC) exhibit enhanced target binding, broadened target capacity, and robust target cleavage compared to invertebrate *Ef*Piwi. MILI and HILI piRISC structures overall resemble the invertebrate *Ef*Piwi and Siwi piRISCs, but we identify distinct features that help to explain biochemical differences between them. Our study helps to explain how MILI and HILI manage to surveil a vast transcriptome using a limited number of piRNAs.

## RESULTS

### MILI and HILI piRISCs bind targets with higher affinity than *Ef*Piwi piRISC

To study how human piRISCs recognize and cleave their targets, we sought to prepare well-behaved piRISC complexes. Exploiting different eukaryotic expression systems, we found that of the four human PIWI proteins—HILI, HIWI, HIWI2, and HIWI3—only HILI could be expressed in HEK 293F cells, albeit still at low levels. HILI is phylogenetically related to Ago3-like PIWI proteins, including *Ephydatia fluviatilis* Piwi (*Ef*Piwi; **Extended Data Fig. 2h**), whose cryo-EM structure has recently been solved^23^. To understand how mammalian piRISC regulates targets, therefore, we expressed HILI or its mouse ortholog MILI and prepared homogenous PIWI-piRNA complexes bound to a synthetic piRNA using an oligo-capture approach ^24^ (**Extended Data Fig. 2a-g**).

To characterize the targeting capability of MILI and HILI piRISC complexes, we measured the equilibrium dissociation constant (*K*_D_) and dissociation rate constant (*k*_off_). MILI and HILI piRISCs bound a seed-complementary target with 6-fold higher affinity than that of *Ef*Piwi piRISC (**Fig. 1a-c**), possibly due to lower target off-rate constants (**Fig. 1b, d**). Comparing to *Ef*Piwi, extended base-pairing through the central region (g9-g14) allowed MILI or HILI to lower the rate constants of target release (**Fig. 1e, Extended Data Fig. 3**). For example, for MILI and HILI, target release rate constants (*k*_off_) decreased to below 0.42×10^-4^ s^-1^ (halftime, t_1/2_ > 4.6h), but 1.33×10^-4^ s^-1^ for *Ef*Piwi (halftime, t_1/2_ > 1.4h). To further compare the binding affinity among MILI, HILI, and *Ef*Piwi, we performed EMSA assay with an entirely complementary target and found that MILI and HILI outperformed *Ef*Piwi (**Fig. 1f**). A different guide RNA and its fully complementary targets (39, 48 or 55 nt) behaved similarly (**Extended Data Fig. 4e**). These results demonstrate that MILI and HILI more effectively recognize targets.

**Fig. 1.**
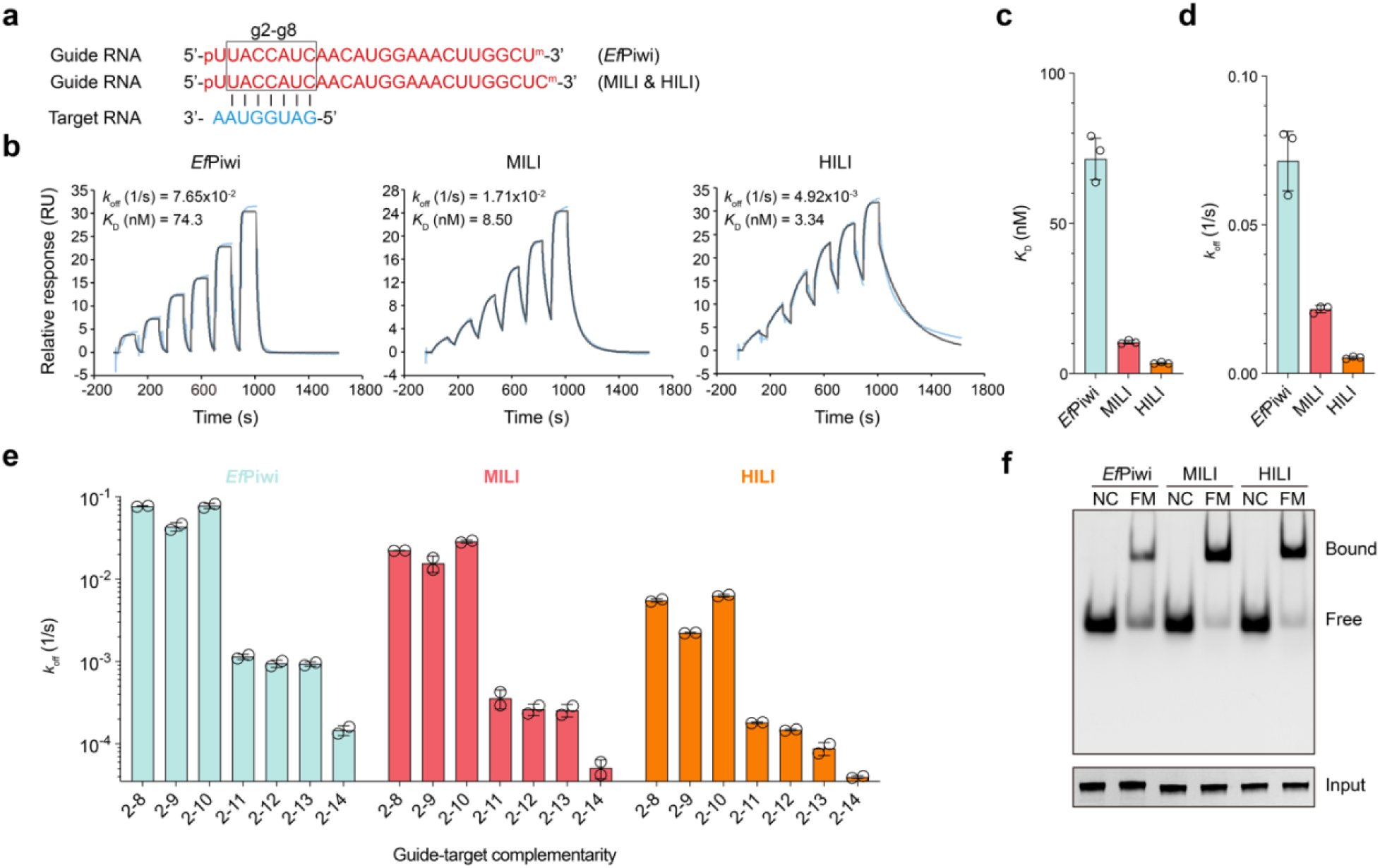
piRNA target binding is more efficient in mammals than in invertebrates. **a**, Guide (red) and target (blue) RNAs used in this experiment. The seed region of the guide (g2– g8) is boxed. **b,** Representative SPR sensorgrams showing binding kinetics of piRISCs to the seed complementary target. Blue trace shows the actual data, and the grey trace shows the binding model fit to the data. **c,** Equilibrium dissociation constant (*K*_D_) of g2–g8 complementary target for protein-guide RNA complexes. **d,** Dissociation rate constant (*k*_off_) of g2–g8 complementary target from protein-guide RNA complexes. **e,** Dissociation rate constant (*k*_off_) of Piwi-piRNA complexes binding target RNAs with various degrees of guide complementarity. **f,** Representative PAGE of the protein-guide RNA complexes binding to fully matched (FM) target RNA. Non-complementary (NC) target of the guide as a negative control. Proteins used in this assay were assessed by silver staining and are shown in the lower panel. In **c-e**, data are expressed as mean ± s.d.

### A wider nucleic-acid-binding channel in PIWI proteins correlates with increased target binding affinity

To better understand the functional differences in seed binding between MILI, HILI, and *Ef*Piwi, we determined the cryo-EM structures of homogeneous MILI and HILI piRISCs at 3.0 and 3.4 Å resolutions, respectively (**Fig. 2a-d, Extended Data Fig. 5a, Extended Data Fig. 6**). The overall domain organization of MILI and HILI piRISC structures, with piRNA threaded through the channel between N-PAZ and MID-PIWI lobes and its 5′ end bound in the MID-PIWI domain and its 3′ end bound in the PAZ domain, resembles that of the silkworm Siwi and sponge *Ef*Piwi piRISC structures^21, 23^. Indeed, as expected, the individual domains of MILI and HILI superimpose well with their counterparts from Siwi and *Ef*Piwi (**Extended Data Fig. 7**), illustrating the overall structural conservation in PIWI-clade Argonautes.

**Fig. 2.**
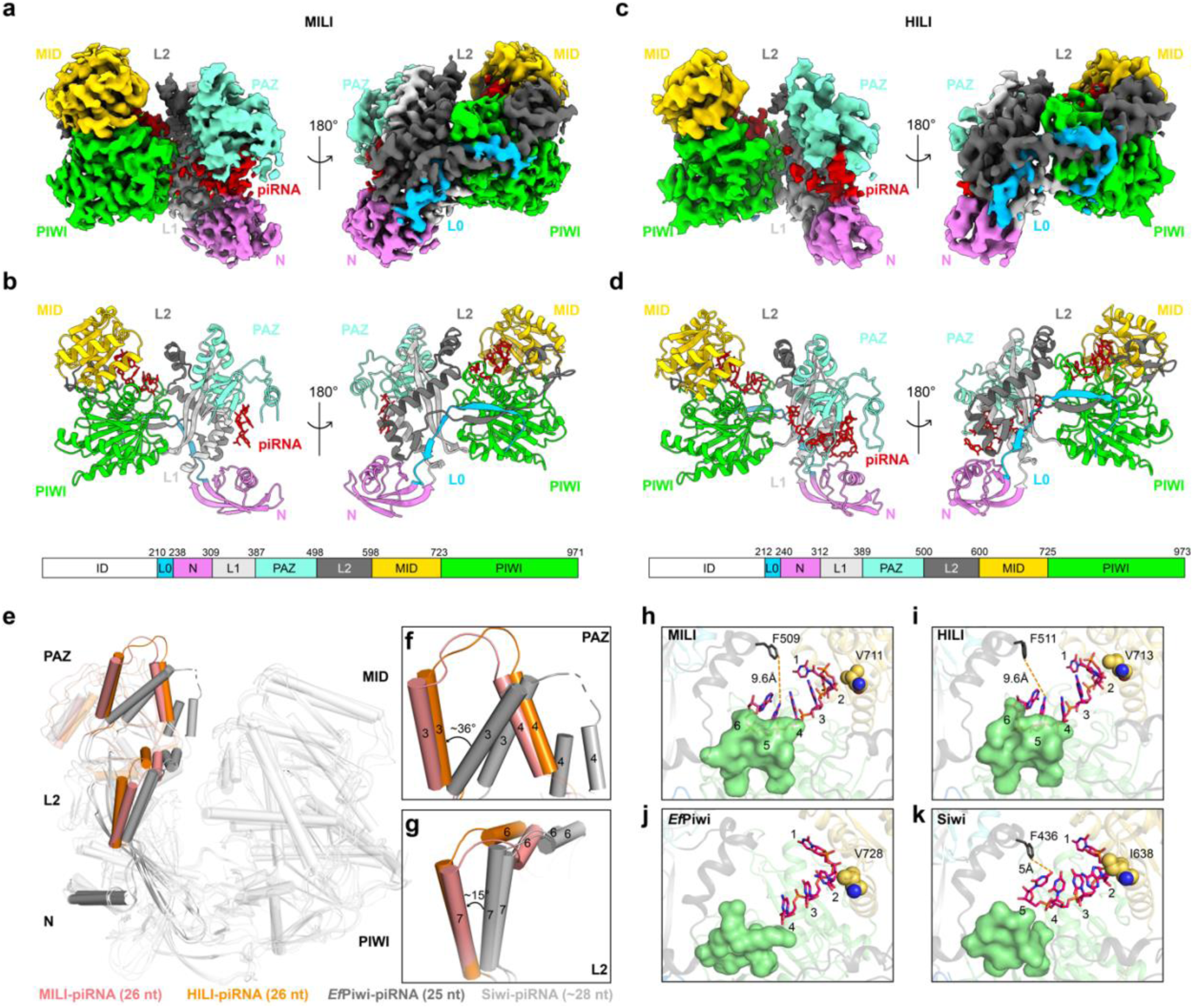
Structure features of mammalian PIWI proteins. **a**, Cryo-EM reconstruction of MILI-piRNA binary complex (colored by domain as indicated). **b,** Cartoon representation of MILI-piRNA. Domain organization is shown below. ID, intrinsically disordered region. **c,** Cryo-EM reconstruction of HILI-piRNA binary complex (colored by domain as indicated). **d,** Cartoon representation of HILI-piRNA. Domain organization is shown below. ID, intrinsically disordered region. **e-g,** Superposition of MILI-piRNA and HILI-piRNA structures and published invertebrate *Ef*Piwi-piRNA (PDB: 7KX7) and Siwi-piRNA (PDB: 5GUH) structures. Structural differences in their PAZ domain (**f**) and seed gate (**g**) are highlighted. In **f**, helices α3 and α4 in PAZ domains of MILI-piRNA and HILI-piRNA shifted away from those of *Ef*Piwi-piRNA and Siwi-piRNA complexes. The approximate angle is indicated. In **g**, helices α6 and α7 (seed gate) of MILI and HILI move outward about 15° relative to their invertebrate counterparts. **h-k,** Seed arrangement of MILI-piRNA, HILI-piRNA, and invertebrate Piwi-piRNA complexes. A conserved phenylalanine residue in helix α7 is shown as sticks in MILI, HILI and Siwi models. g5–g6 nucleotide-binding loops that may result in steric clashes are shown as surface in green.

Notably, we also observed unique structural features that could explain why mammalian piRISCs bind targets with higher affinity than their invertebrate counterparts. First, the nucleic acid-binding channel from MILI or HILI is wider than from Siwi or *Ef*Piwi. The channel is widened by a 36° rotation of PAZ-domain helices α3 and α4 away from the MID-PIWI lobe and a 15° rotation of L2-linker helices α6 and α7 away from MID-PIWI (**Fig. 2e–g**). Second, the seed region of the piRNA guide adopts an A-form conformation in both MILI and HILI piRISCs, but not in Siwi or *Ef*Piwi piRISCs. Indeed, steric hindrance in the vicinity of piRNA nucleotides g5-g6 in Siwi and *Ef*Piwi piRISCs prevent base-stacking interactions needed to adopt an A-form helix (**Fig. 2j-k**)^21, 23^. In Siwi, for example, a conserved phenylalanine (Phe436) from L2-domain helix 7 interacts with g5 and prevents g5-g6 base stacking **(Fig. 2k)**. In MILI and HILI, however, the relative outward shift of L2-domain helix 7 positions the conserved phenylalanine (Phe509 in MILI and Phe511 in HILI) ∼9.6 Å from g5 (**Fig. 2h-i, Extended Data Fig. 8**). These results suggest that the wider nucleic-acid-binding channel in MILI and HILI piRISCs allows the seed region to adopt an A-form conformation, which is presumed to minimize the entropic cost and initiate stable piRNA–target duplex formation, similar to miRNA targeting^25–27, 28–30^. Indeed, the seed-binding affinities of MILI and HILI piRISCs were comparable to that of Ago2 miRISC (**Extended Data Fig. 4a-d**). These observations suggest that the slightly wider nucleic-acid binding-channel in MILI and HILI piRISCs allows them to initiate more efficient target binding than *Ef*Piwi piRISC.

### MILI and HILI piRISCs tolerate seed mismatches

We next determined a 3.4-Å resolution cryo-EM structure of MILI-piRISC bound to a fully complementary 15-nucleotide target RNA (**Fig. 3a, Extended Data Fig. 5b**). The piRNA-target RNA duplex (spanning positions g2 to g15) adopts an A-form double helix, accommodated by a 25° shift of the N-PAZ lobe away from the MID-PIWI lobe (**Fig. 3b**). In this open conformation, the seed gate is displaced by ∼11° compared to *Ef*Piwi-piRNA-target (16nt) complex and does not contact the minor groove of the piRNA–target duplex (**Fig. 3d**). By contrast, in the *Ef*Piwi-piRNA-target ternary complex^23^, the seed gate docks with the minor groove of piRNA–target duplex near the 3′ end of the seed, which enforces more faithful seed pairing *via* extensive van der Waals and hydrophobic interactions from residues R525, M531, K532, and R539 (**Fig. 3c**).

**Fig. 3.**
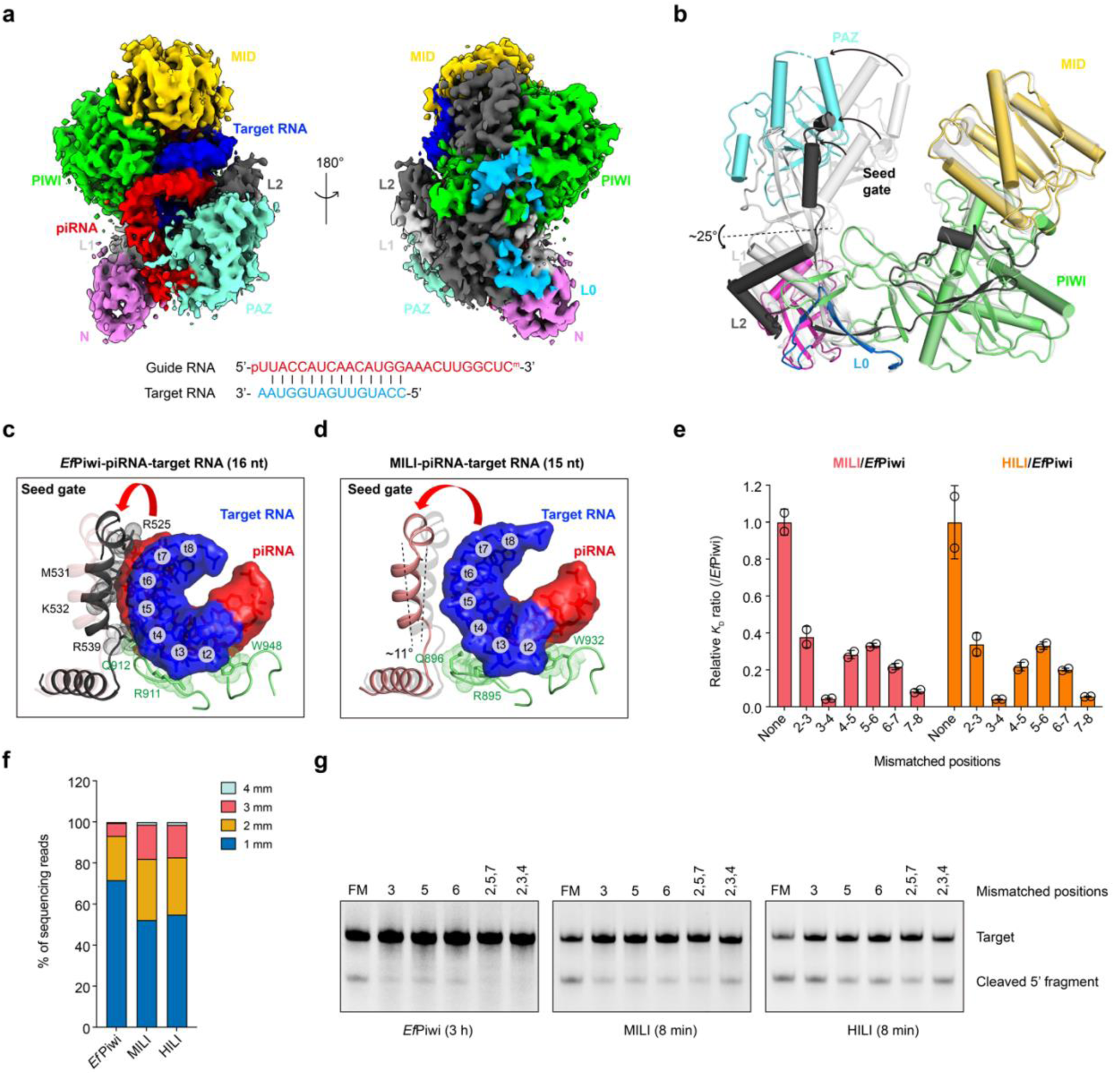
Structural basis for piRNA targeting. **a,** Cryo-EM reconstruction of MILI-piRNA-target RNA (15 nt) ternary complex (domains colored as in Fig. 2). Sequence of the piRNA–target RNA duplex is shown below. **b,** Superposition of MILI-piRNA (white gray, semi-transparent) and MILI-piRNA-target RNA (15 nt) (colored, solid) structures. Arrows indicate movement from guide-only to guide-target structures. Dashed line marks the hinge in the L1 domain. **c,** Interactions between *Ef*Piwi protein and piRNA-target RNA (16 nt) duplex around the seed. **d,** Interactions between MILI protein and piRNA-target RNA (15 nt) duplex around the seed. **e,** The relative *K*_D_ ratio of MILI or HILI over *Ef*Piwi. Relative ratio is calculated as *K*_D_ (mm)/*K*_D_ (none) of MILI or HILI divided by *K*_D_ (mm)/*K*_D_ (none) of *Ef*Piwi. none: no mismatched target, mm: mismatched target, *K*_D_: equilibrium dissociation constant. Plotted data are represented as mean ± s.d. from at least two independent experiments. **f,** Percentage of RNA sequencing reads of cleavage products. Total reads are normalized to 1,000,000 for each sample. **g,** Representative urea-PAGE showing *Ef*Piwi-, MILI-, or HILI-mediated cleavage of targets with 0, 1, 3 mismatches.

Displacement of the seed gate from the piRNA–target duplex suggested that MILI might tolerate base-pair mismatches in the piRNA seed region. We therefore measured the binding affinities of MILI, HILI, and *Ef*Piwi to target RNAs with dinucleotide mismatches to the seed (**Extended Data Fig. 9**). Seed mismatches reduced the binding affinities of all three piRISCs, but the binding affinity of *Ef*Piwi was more reduced than that of MILI or HILI (**Fig. 3e**). For example, a dinucleotide mismatch at g3-g4 increased the dissociation constant (*K*_D_) of all three piRISCs, but the dissociation constant (*K*_D_) of *Ef*Piwi piRISC was 20-fold higher than MILI or HILI piRISCs. These data suggest that the open seed gate of mammalian piRISCs enables better tolerance of seed base-pairing mismatches.

To measure the effect of seed mismatches on target cleavage, we incubated excess MILI, HILI, or *Ef*Piwi piRISCs with a pool of target RNAs comprising equimolar amounts of 63 targets with 1, 2, 3, or 4 mismatches to the piRNA seed sequence (**Extended Data Table 2**). We then analyzed the cleavage products by deep sequencing. As expected, the ability of piRISC to cleave a target decreased as the number of seed mismatches increased, with less than 2% of cleavage products derived from targets with 4 mismatches (**Fig. 3f**). Compared to *Ef*Piwi piRISC, however, MILI and HILI piRISCs generated 2- to 3-times as many cleavage products from targets with 3 or 4 mismatches. We observed similar results in gel-based analyses of cleavage assays using targets with 0, 1, or 3 mismatches (**Fig. 3g**). Note that extended incubation was required to observe cleavage products generated by *Ef*Piwi piRISC, consistent with previous studies showing that *Ef*Piwi and Siwi are feeble endonucleases^23^. These results suggest that the MILI and HILI piRISCs tolerate seed mismatches for target cleavage.

Outside of the seed-paired region, we observed that MILI—like *Ef*Piwi—recognizes the overall helical structure of the piRNA-target duplex *via* contacts with the sugar-phosphate backbone ^23^ (**Extended Data Fig. 10a-b**), suggesting that MILI tolerates mismatches beyond the seed. Indeed, trinucleotide mispairing from g11 to g22 had little effect on target binding by *Ef*Piwi, MILI, or HILI piRISCs (**Extended Data Fig. 10c**). Taken together, the tolerance of piRNA-target mismatches in seed and non-seed regions is expected to broaden the targeting potential of mammalian piRNAs, making them well-suited to adapt to ever-changing genomic threats.

### MILI and HILI piRISCs are inherently more robust endonucleases than *Ef*Piwi piRISC

In cleavage assays with fully matched target RNAs, MILI, and HILI piRISCs generated 25-fold more cleavage product than did *Ef*Piwi piRISC (**Fig. 4a, Extended Data Fig. 11a**). MILI and HILI remained more active than *Ef*Piwi and Siwi in the presence of the auxiliary protein GTSF1^31^, which enhanced the cleavage activities of MILI, HILI, *Ef*Piwi, and Siwi (**Extended Data Fig. 12**). In addition, Siwi was similar to *Ef*Piwi in target binding, cleavage activity, and mismatch tolerance (**Extended Data Fig. 13)**. The inherently higher cleavage activities of MILI and HILI piRISCs might result from increased target binding affinity, higher catalytic activity, or more rapid release of products to regenerate active piRISC. Indeed, MILI and HILI piRISCs bound with higher affinity to fully complementary target RNA of different lengths than did *Ef*Piwi piRISC (**Fig. 4b**). To measure the initial target cleavage rate (*V*_0_), we incubated piRISCs in vast excess with perfectly complementary target RNA molecules—i.e., [E]>>[S]—such that piRISCs can only bind and cleave one target RNA^32^. We found that MILI and HILI piRISCs cleaved at an initial velocity of 0.658 nM min^-1^ and 0.904 nM min^-1^, respectively, at least 25-fold greater than that of *Ef*Piwi piRISC (0.025 nM min^-1^; **Fig. 4c, Extended Data Fig. 11b**). Moreover, the catalytic constants (*k*_cat_) of MILI and HILI piRISCs were at least 20-fold greater than that of *Ef*Piwi (**Fig. 4d**). Lastly, MILI, HILI, and *Ef*Piwi piRISCs released cleavage products with similar slow kinetics (**Fig. 4e, Extended Data Fig. 11c**). Together, these results indicate that MILI and HILI piRISCs bind and cleave substrates more efficiently than *Ef*Piwi piRISC.

**Fig. 4.**
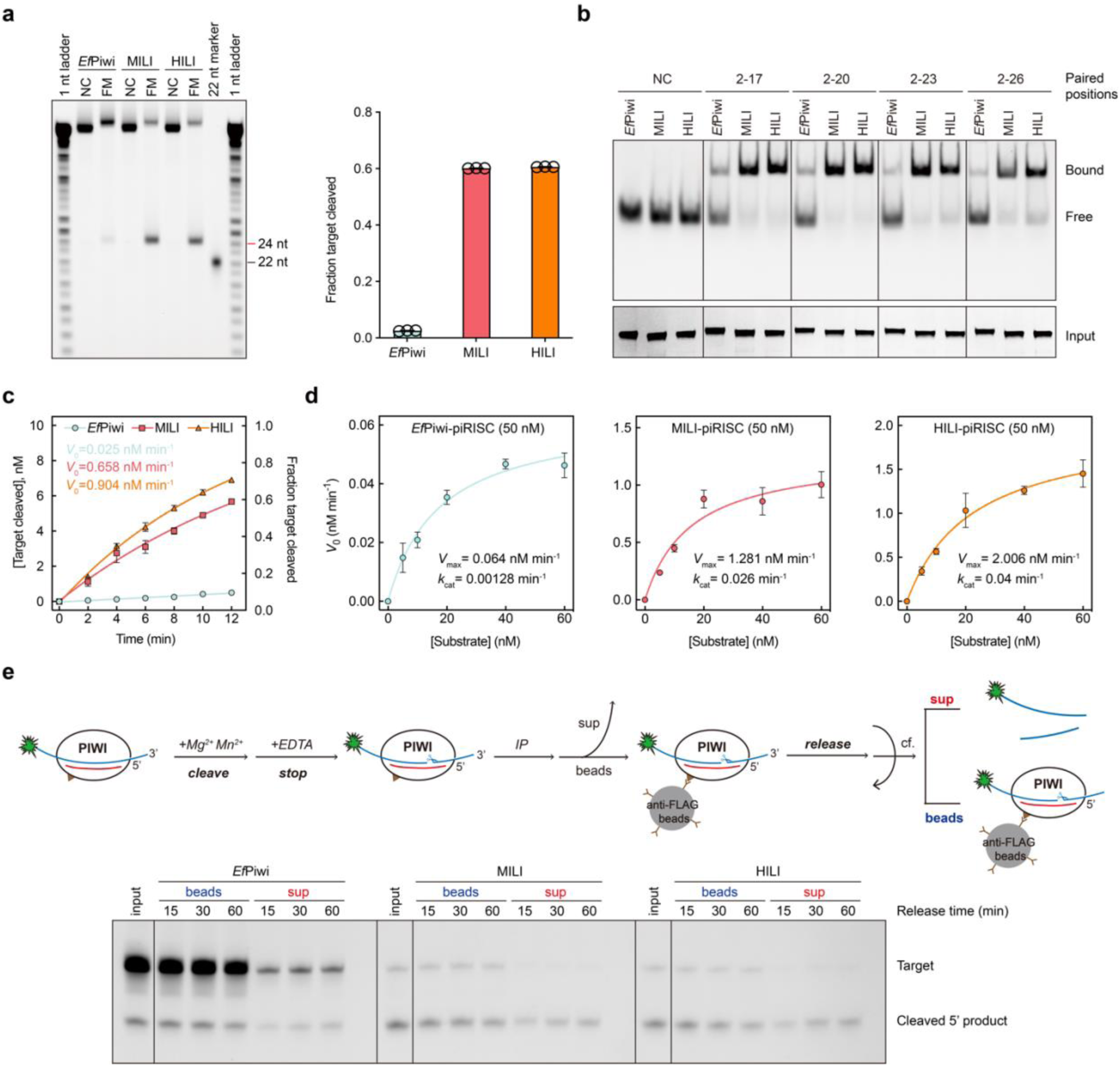
Biochemical characterization of target cleavage by MILI and HILI. **a**, Representative *in vitro* cleavage reactions with equimolar amounts of substrate (target RNA) and enzyme (Piwi-piRNA complexes). Quantification of the *in vitro* cleavage assays (right). NC, non-complementary target as negative control. Data are represented as mean ± s.d. from independent triplicates. **b,** Representative native PAGE of the protein-guide RNA complexes binding to target RNA with complementarity to nucleotides g2–g17, g2–g20, g2–g23, and g2–g26. Proteins used in this assay were assessed by silver staining and are shown in the lower panel. **c,** Target cleavage assays in presence of excess enzyme, i.e., [S] (target RNA) < [E] (Piwi-piRNA complexes). Initial rates, *V*_0_, were determined by fitting the data to a single exponential. Plotted curves are represented as mean ± s.d. from three independent experiments. **d,** Kinetics of purified *Ef*Piwi-piRISC, MILI-piRISC and HILI-piRISC with a 26-nt fully complementary target. Data are mean ± s.d. for three independent experiments fitted to Michaelis-Menten. **e,** Flow chart of target release assay upon cleavage. Different amounts of Flag-tagged Piwi-piRNA complexes were incubated with 5′-FAM-labeled target RNA in the presence of divalent cations to accumulate cleavage product, followed by the addition of EDTA to stop the reaction. After immunoprecipitation (IP) by anti-Flag beads, supernatant (sup) was discarded and bound cleaved target RNAs in the beads were deemed as total cleaved target RNAs (input). Beads were then suspended in cleavage reaction buffer without MgOAc and MnCl_2_ and incubated for 15, 30, or 60 mins at 37°C. Input, sup, and beads fractions were analyzed by denaturing urea-PAGE.

### A vertebrate-specific residue interacts with the piRNA seed to lower the activation threshold of MILI and HILI

Compared to the *Ef*Piwi ternary complex, the phosphate backbone of the piRNA-target duplex in MILI is locally distorted at positions g5 and g6 (**Fig. 5a**), with increasing displacements moving towards the distal end (**Fig. 5b**). As a result, the distal end of the piRNA-target duplex is closer to the PAZ domain in the MILI ternary complex, possibly in favor of outside movement of the PAZ domain and the expansion of the N-PAZ channel, which is necessary for target cleavage activation in RISC^28–30^. Notably, Lys943 of MILI protrudes from helix α19 of the PIWI domain and forms hydrogen bonds with phosphate 5 of the piRNA (**Fig. 5c**). This lysine is conserved in Ago3-like vertebrate PIWI proteins (e.g., K945 in HILI), but not in invertebrate PIWI proteins (**Fig. 5c-d, Extended Data Fig. 14**). In *Ef*Piwi, for example, the corresponding position is Asn959, which does not reach the phosphate backbone of the piRNA guide (**Fig. 5c-d**). We hypothesized that the conserved lysine in α19 confers superior target binding and cleavage activity on MILI and HILI piRISCs compared to *Ef*Piwi piRISC. If correct, we reasoned that mutating Asn959 to Lys (N959K) would increase the target RNA binding and cleavage activity of *Ef*Piwi piRISC. Conversely, mutating MILI Lys943 or HILI Lys945 to Asn should reduce the target RNA binding and cleavage activity of MILI and HILI piRISCs. Indeed, *Ef*Piwi^N959K^ piRISC showed increased target binding and cleavage activities compared to wild-type *Ef*Piwi piRISC (**Fig. 5e-f**). By contrast, MILI^K943N^ and HILI^K945N^ mutants showed reduced target binding and cleavage activities compared to wild-type MILI and HILI piRISCs (**Fig. 5g-h**). Thus, this conserved lysine potentiates target binding and cleavage by vertebrate Ago3-like PIWIs.

**Fig. 5.**
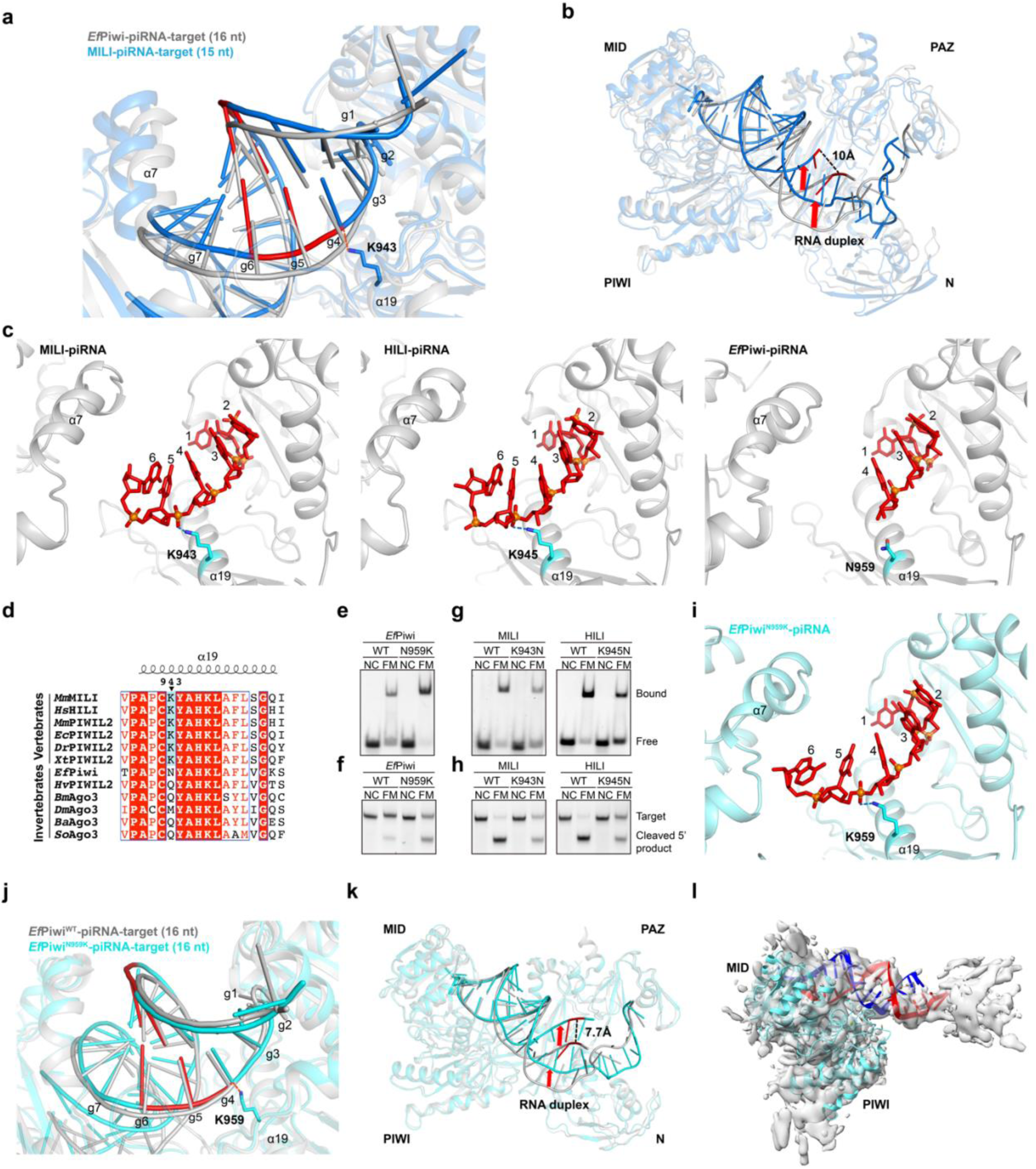
Mutation analysis of mammalian-specific amino acid residue. **a**, Close-up view of the seed region of superimposed *Ef*Piwi-piRNA-target (16 nt) and MILI-piRNA-target (15 nt) complexes. **b,** Overall view of superimposed *Ef*Piwi-piRNA-target (16 nt) and MILI-piRNA-target (15 nt) complexes. Red arrows indicate the shifted trajectory of the guide-target duplex in the MILI-piRNA-target (15 nt) compared to that in the *Ef*Piwi-piRNA-target (16 nt) complex. **c,** Comparison of the seed regions showing K943 in MILI-piRNA, K945 in HILI-piRNA, and N959 in *Ef*Piwi-piRNA. The guide strand is shown in red, with phosphorus atoms as orange balls. **d,** Sequence alignment of ⍺19 region of vertebrate and invertebrate Ago3-like PIWI proteins, showing that Lys943 of MILI is a vertebrate-specific Ago3-like PIWI proteins. Species abbreviations are defined in the legend of Extended Data Fig. 14. **e,f,** Effect of N959K mutation on *Ef*Piwi-piRNA target binding (**e**) and cleavage (**f**). **g,h,** Effects of K943N mutation (MILI-piRNA) and K945N (HILI-piRNA) on target binding (**g**) and cleavage (**h**). **i,** Structure of region surrounding K959 (*Ef*Piwi^N959K^-piRNA). **j,** Close-up view of the seed region of superimposed *Ef*Piwi^WT^-piRNA-target (16 nt) and *Ef*Piwi^N959K^-piRNA-target (16 nt) complexes. **k,** Overall view of superimposed *Ef*Piwi-piRNA-target (16 nt) and *Ef*Piwi^N959K^-piRNA-target (16 nt) complexes. Red arrows indicate the shifted trajectory of the guide-target duplex in the *Ef*Piwi^N959K^-piRNA-target (16 nt) complex away from that in the *Ef*Piwi-piRNA-target (16 nt) complex. **l,** Cryo-EM reconstruction of “open” *Ef*Piwi^N959K^-piRNA-target (16 nt) complex. The guide strand is shown in red and target strand is in blue.

Lastly, we solved a 3.2-Å cryo-EM structure of an *Ef*Piwi^N959K^-piRNA complex and a 3.2-Å structure of an *Ef*Piwi^N959K^-piRNA-target (15-nt) ternary complex (**Extended Data Fig. 15**). In the absence of target, *Ef*Piwi^N959K^ piRISC displayed an extended piRNA seed, resembling that observed in MILI and HILI piRISCs (**Fig. 5i, Extended Data Fig. 15a**). Notably, the *Ef*Piwi^N959K^-piRNA-target ternary features a piRNA target duplex is locally distorted at g5 and increasingly displaced towards the distal end compared to that in *Ef*Piwi^WT^-piRNA-target ternary complex (**Fig. 5j-k**). For example, the target nucleotide t15 in the *Ef*Piwi^N959K^-piRNA-target ternary complex is shifted nearly 7.7 Å from that in *Ef*Piwi^WT^-piRNA-target ternary complex, similar to the shift observed in MILI compared to *Ef*Piwi^WT^ (10 Å; **Fig. 5b**).

Previous work showed that extensive piRNA–target pairing stimulates *Ef*Piwi nuclease activity. In our hands, *Ef*Piwi nuclease activity required a minimum 19 base-pair complementarity (g2–g20). MILI and HILI nuclease activities were also stimulated by extended piRNA–target pairing, but both MILI and HILI cleaved targets with only 14 base-pair complementarity (g2–g15) (**Extended Data 16c-e**). Moreover, the fraction of cleaved targets generated *in vitro* by MILI or HILI with g2–g15 pairing was 3-fold higher than the maximal level of cleavage by *Ef*Piwi (**Extended Data Fig. 16d**). An analysis of cleaved piRNA targets in mouse primary spermatocytes similarly shows that mouse PIWIs (MIWI and MILI) cleave targets complementary to g2–g13 and that extended piRNA–target pairing increases the fraction of targets cleaved *in vivo*^33^. By contrast, a similar analysis of Ago3-dependent degradome data from *Drosophila* ovaries^34, 35^ suggests that *Drosophila* Ago3 nuclease activity may require 21 base-pair complementarity and generates fewer cleavage products per target *in vivo* than mouse PIWIs (fraction cleaved, 0.048 for Ago3 g2–g24 vs. 0.52 for MILI/MIWI g2–g20)^33^ (**Extended Data Fig. 16f-h**).

Notably, *Ef*Piwi piRISC bound to a fully complementary target (from g2–g25) yielded cryo-EM particles in which the piRNA–target duplex extends from the MID-PIWI lobe but is uncoupled from the N-L1-PAZ-L2 lobe, suggesting that extended piRNA–target base pairing activates cleavage by widening the central cleft^23^. In our cryo-EM data, this open possibly “activated” state comprises 59.3% of *Ef*Piwi^N959K^ particles and 29.6% of MILI particles (**Fig. 5l, Extended Data Fig. 5b, Extended Data Fig. 15b**). Together, these observations suggest that the conserved lysine on α19 of vertebrate PIWI (e.g., Lys943 of MILI) plays a key role in promoting the function of MILI and other vertebrate homologs.

## DISCUSSION

Structural studies of miRNA-mediated silencing suggest that Argonaute proteins use their small RNA guides to recognize their targets *via* a stepwise process initiated by seed pairing, which in turn widens the nucleic-acid-binding channel to allow subsequent base pairing and stimulate target cleavage^28–30, 36^. Our findings suggest that vertebrate Ago3-like MILI and HILI piRISCs adopt a slightly open nucleic-acid-binding channel with an extended seed that improves the rate and affinity of target binding as well as cleavage activity compared to invertebrate Ago3-like piRISCs. Target binding widens the channel further by shifting the seed gate and elements of the PAZ domain to accommodate the propagation of guide–target pairing. Moreover, a single lysine residue—conserved in vertebrate Ago3-like PIWI proteins—interacts with the phosphate backbone between g4 and g5 of the piRNA guide, which shifts the trajectory of the piRNA–target duplex out of the nucleic-acid-binding channel and toward the PAZ domain. This conformation is consistent with the recently proposed model for *Ef*Piwi activation, whereby extended piRNA– target pairing forces the PAZ domain to uncouple from the PIWI domain, opening the central cleft and activating *Ef*Piwi. Our findings suggest that a vertebrate-specific lysine lowers the activation threshold of MILI and HILI piRISCs compared to *Ef*Piwi piRISC. Further investigation is needed to determine how other conformational differences or auxiliary factors account for differences between MILI/HILI and invertebrate piRISC function *in vitro* or *in vivo*. For example, recent studies suggest that VASA-like RNA helicases promote target release from Argonaute proteins^35, 37^.

Effective defense against nucleic acid pathogens might be established through potent cleavage and swift turnover of cleaved targets and ping-pong piRNA amplification fueled by target cleavage^38^. Remarkably, recent work has shown that mouse PIWIs tolerate a target nucleotide mismatch at any position, including mismatches at the cleavage site^33, 39^. The more open nucleic-acid-binding channel of MILI and HILI piRISCs appears to correlate with an ability to tolerate mismatches even in the seed region. Put another way, the shifted seed gate in MILI and HILI piRISCs mitigates the need to monitor ideal seed pairing. As such, increased tolerance for seed mismatches broadens the targeting capacity of MILI and HILI piRISCs, suppressing the escape of targets. The fidelity of targeting is enforced by extended base pairing into the central region of the piRNA (**Extended Data Fig. 16a-b**), while target cleavage is activated by base pairing toward the piRNA 3’ end (**Extended Data Fig. 16c-e**). Thus, extended base pairing by MILI, HILI, and *Ef*Piwi ensures accurate and efficient regulation of transposons and meanwhile sufficient to prevent inappropriate targeting of mRNA and long non-coding RNAs^33^, even if a transposon target site contains a few mismatches (but seed mismatches are less tolerated by *Ef*Piwi or Siwi). Nevertheless, MILI and HILI piRISCs also support binding and cleavage of targets with as little as 14-nt guide-target complementarity (g2–g15), perhaps enabling them to tune the expression of a subset of mRNAs required for spermatogenesis^40–44^.

Our structural and biochemical characterization of mammalian piRISCs has uncovered critical features that uniquely distinguish vertebrate Ago3-related piRISCs from invertebrate counterparts. Notably, our work reveals the first structure and mechanistic features of a human PIWI protein, thereby helping us decipher the mechanism of human PIWI-mediated genome defense and gene regulation as well as its roles in human fertility and disease^45, 46^.

---

**Extended Data Fig. 1.**
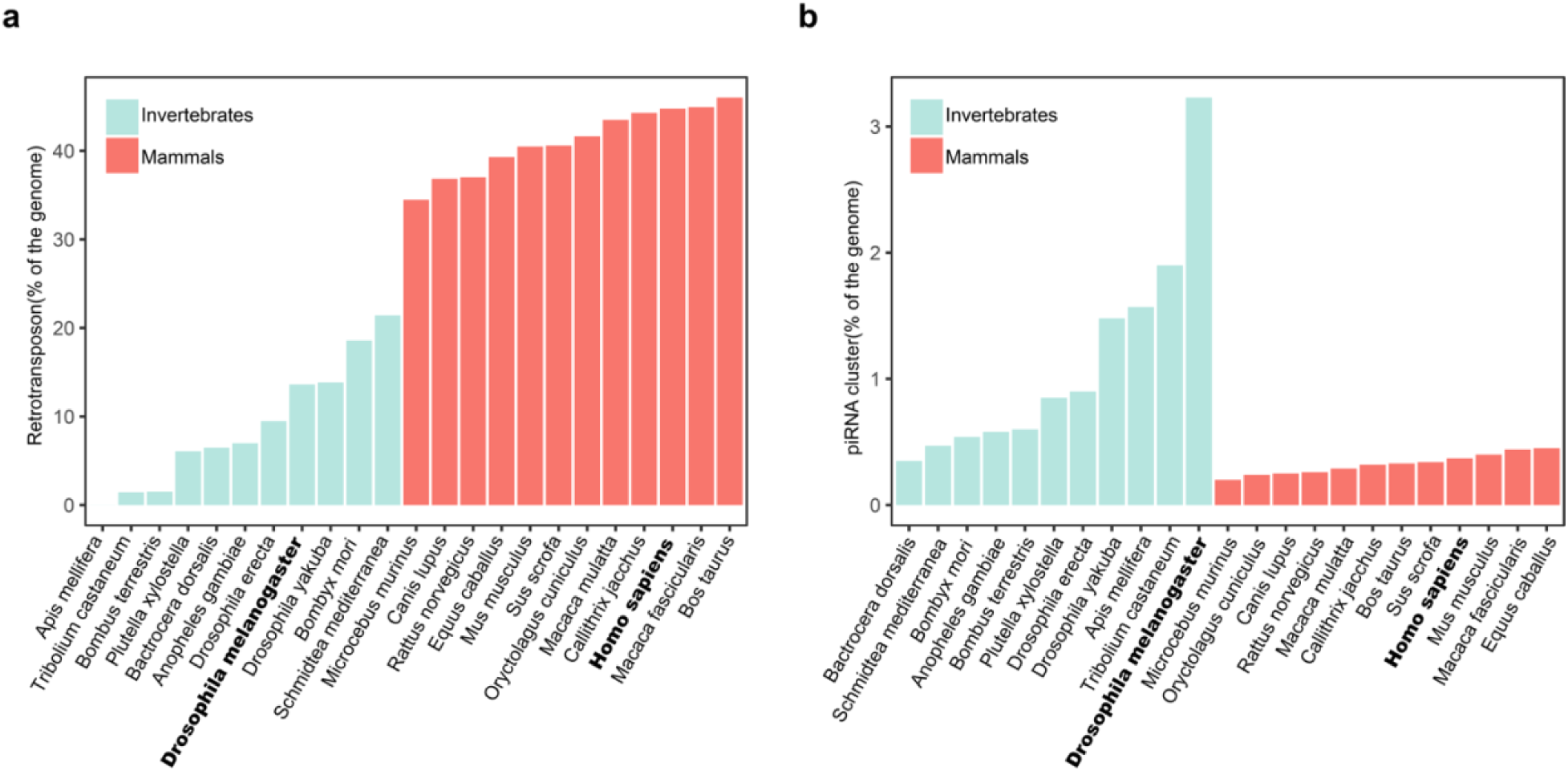
Retrotransposon and piRNA cluster in invertebrates and mammals. **a,b**, Ratio of retrotransposon (**a**) and piRNA cluster (**b**) of invertebrate and mammalian genomes.

**Extended Data Fig. 2.**
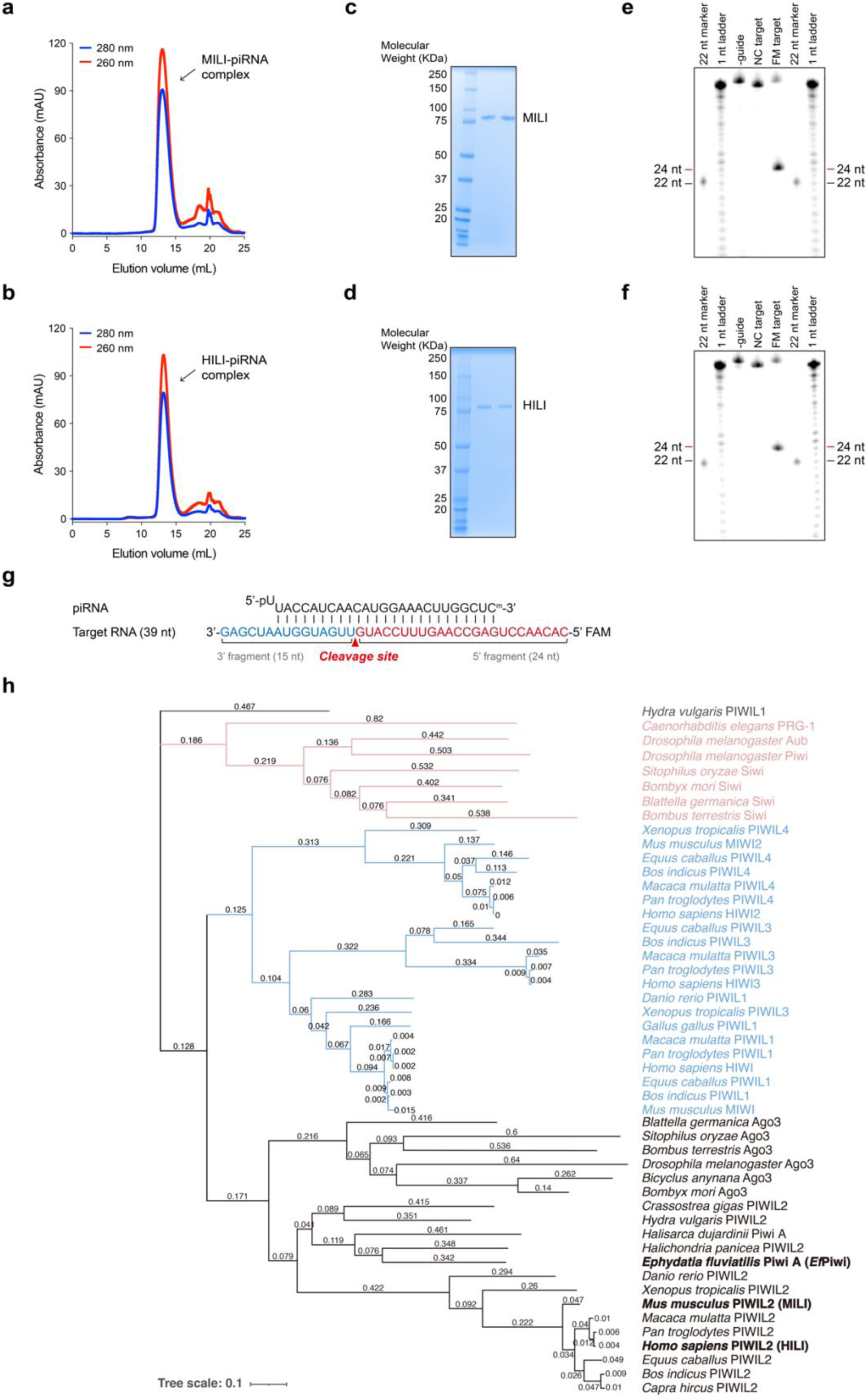
Preparation of Piwi-piRNA complexes and PIWI family tree. **a,b**, Size-exclusion chromatography profile of the purified MILI-piRNA complex (arrow) (**a**) and HILI-piRNA complex (arrow) (**b**) using a modified oligo capture approach. **c,d,** SDS-PAGE analysis of the purified MILI-piRNA complex (**c**) and HILI-piRNA complex (**d**). Gels were stained with Coomassie blue. **e,f,** Guide RNA-mediated cleavage assay with MILI (**e**) or HILI (**f**). MILI or HILI loaded with or without synthetic guide RNAs were incubated with 5′-FAM-labeled fully matched (FM) target RNAs or non-complementary (NC) target RNAs for 1 h at 37°C. Products were resolved on a denaturing urea-PAGE gel alongside synthetic 22-nt marker and 1-nt RNA ladder. **g,** piRNA and target RNA sequences used in **e,f**. The fluorescein amidite (FAM) modification in the target strand is highlighted in green, and the cleavage site is indicated with a red arrow. The cleaved 5′ fragment with FAM label is 24 nucleotides. **h,** Phylogram of the PIWI proteins shows that MILI, HILI, and *Ef*Piwi proteins belong to the same branch. Branch lengths are shown above the line.

**Extended Data Fig. 3.**
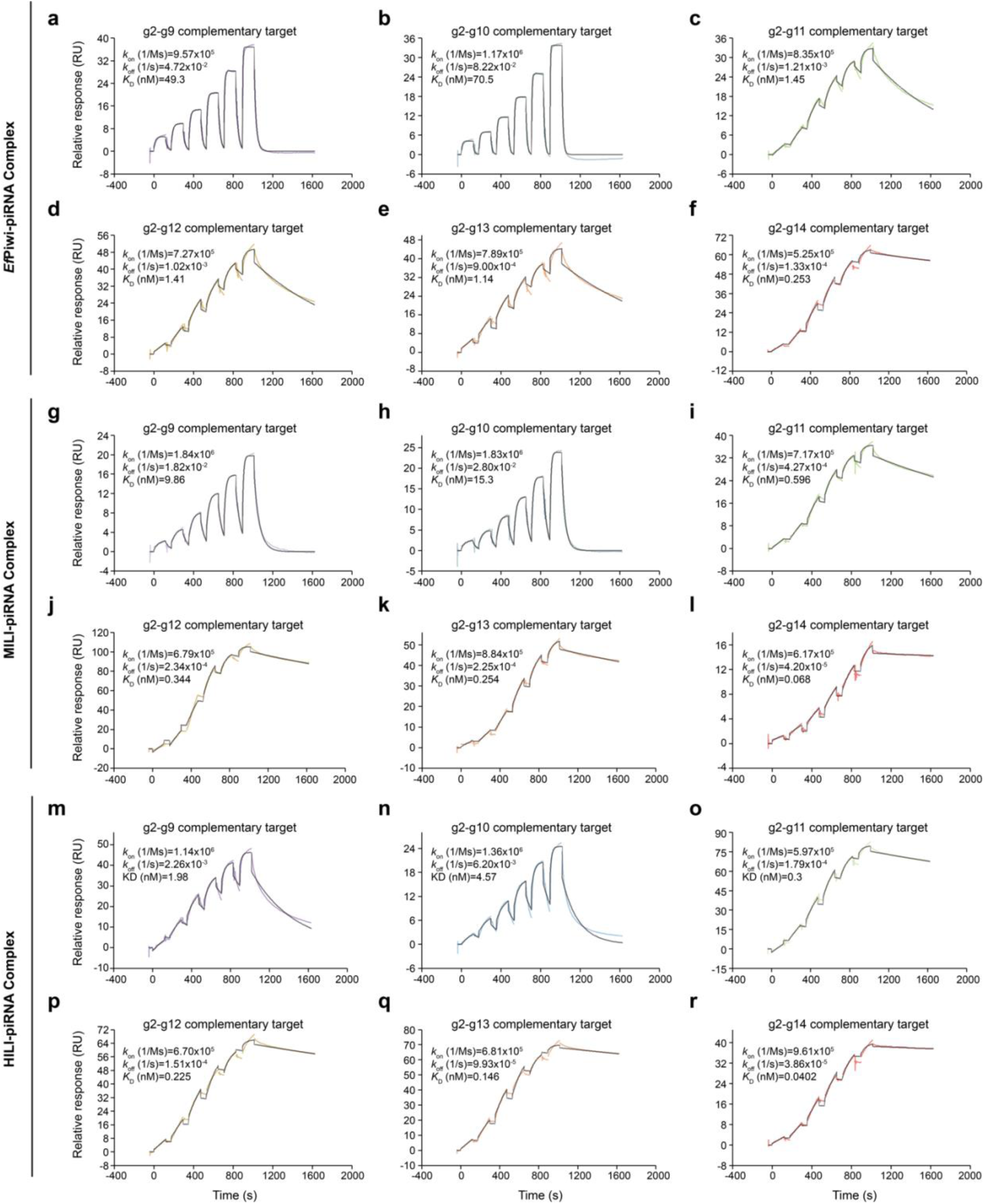
Kinetics of Piwi-piRNA complexes binding target RNAs with various degrees of guide complementarity. **a-f**, Representative binding kinetics of *Ef*Piwi-piRNA complex binding target RNAs with complementarity to nucleotides g2-g9 (**a**), g2-g10 (**b**), g2-g11 (**c**), g2-g12 (**d**), g2-g13 (**e**), g2-g14 (**f**). **g-l,** Representative binding kinetics of MILI-piRNA complex binding target RNAs with complementarity to nucleotides g2-g9 (**g**), g2-g10 (**h**), g2-g11 (**i**), g2-g12 (**j**), g2-g13 (**k**), g2-g14 (**l**). **m-r,** Representative binding kinetics of HILI-piRNA complex binding target RNAs with complementarity to nucleotides g2-g9 (**m**), g2-g10 (**n**), g2-g11 (**o**), g2-g12 (**p**), g2-g13 (**q**), g2-g14 (**r**). Data shown in rainbow colors and fit the binding model shown in the grey line.

**Extended Data Fig. 4.**
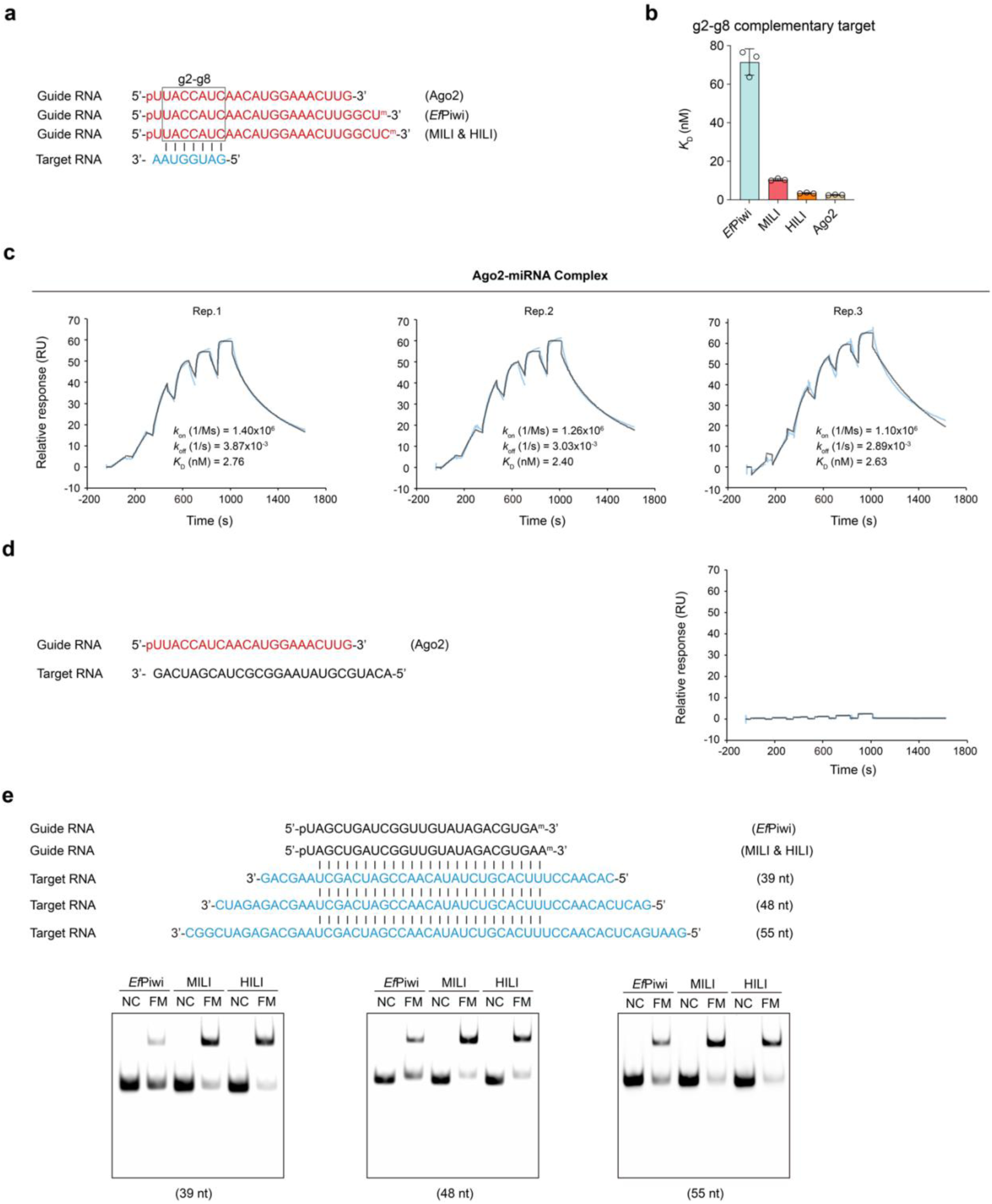
Kinetics of Ago2-miRNA target binding and EMSA of piRISC binding to target RNAs. **a**, The guide and target RNAs used in this experiment. **b,** Equilibrium dissociation constant (*K*_D_) of g2-g8 complementary target for protein-guide RNA complexes. Data are mean ± s.d. from independent triplicates. **c,** SPR traces showing binding kinetics of g2-g8 complementary target for Ago2-miRNA binary complex. **d,** SPR sensorgrams of non-complementary target RNA binding to Ago2-miRNA complex. **e,** Representative PAGE of the protein-guide RNA complexes binding to another fully matched (FM) target RNA. Non-complementary (NC) target is a negative control.

**Extended Data Fig. 5.**
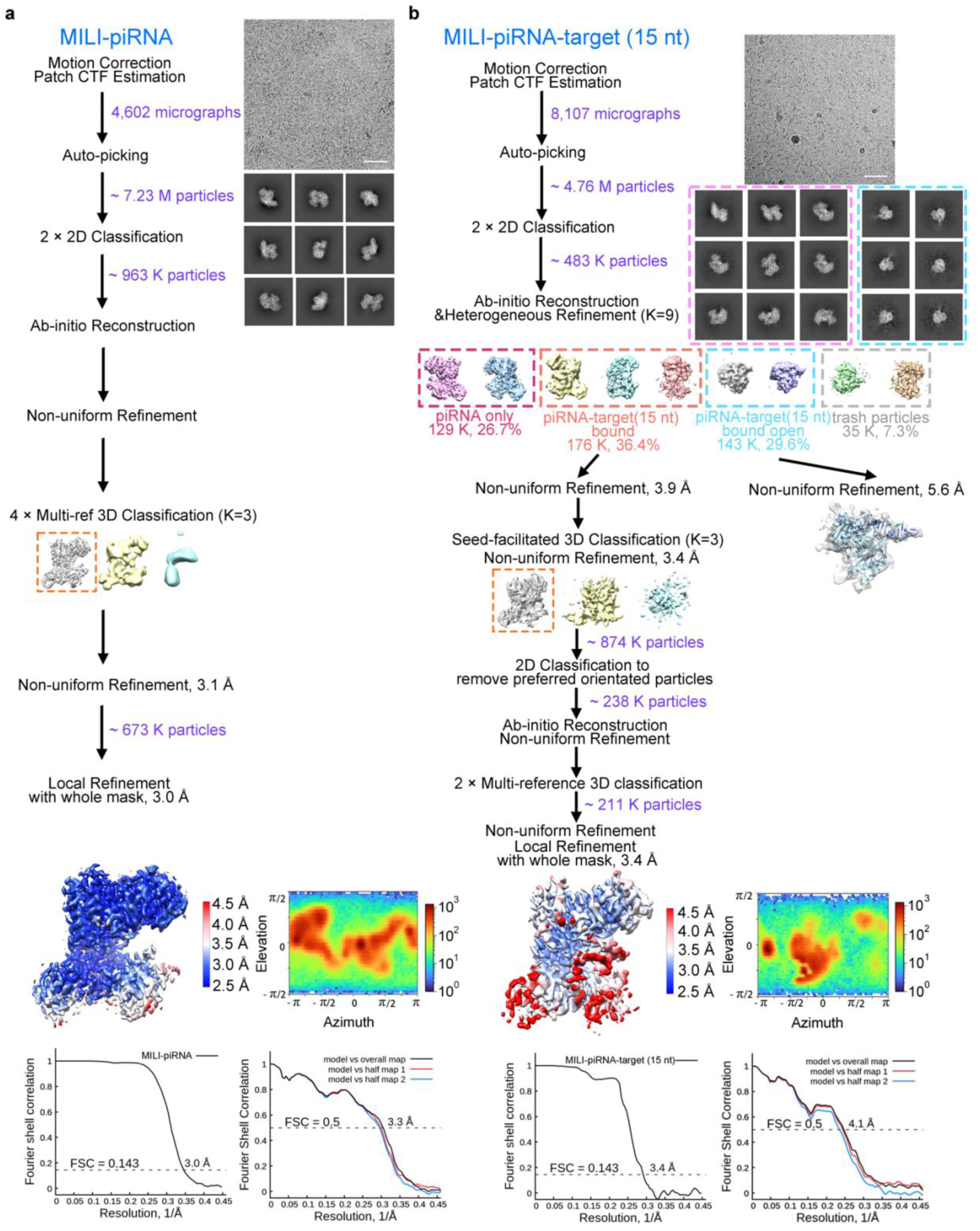
Cryo-EM analysis of MILI-piRNA, MILI-piRNA-target (15 nt). **a,b**, Data processing workflow of cryo-EM analysis of MILI-piRNA (**a**) and MILI-piRNA-target (15 nt) (**b**). Representative motion-corrected dose-weighted micrograph (out of 4,602 and 8,107 micrographs) with a scale bar of 50 nm, 2D class averages with a box size of 209 Å, 3D classification, 3D refinement reconstruction with local resolution, angular distribution, gold-standard FSC curves of the final map (resolution cut-off at FSC = 0.143), and model vs. map curves of the final map against the final model (resolution cut-off at FSC = 0.5). 3D classes selected for subsequent analysis were indicated with dotted boxes in different colors.

**Extended Data Fig. 6.**
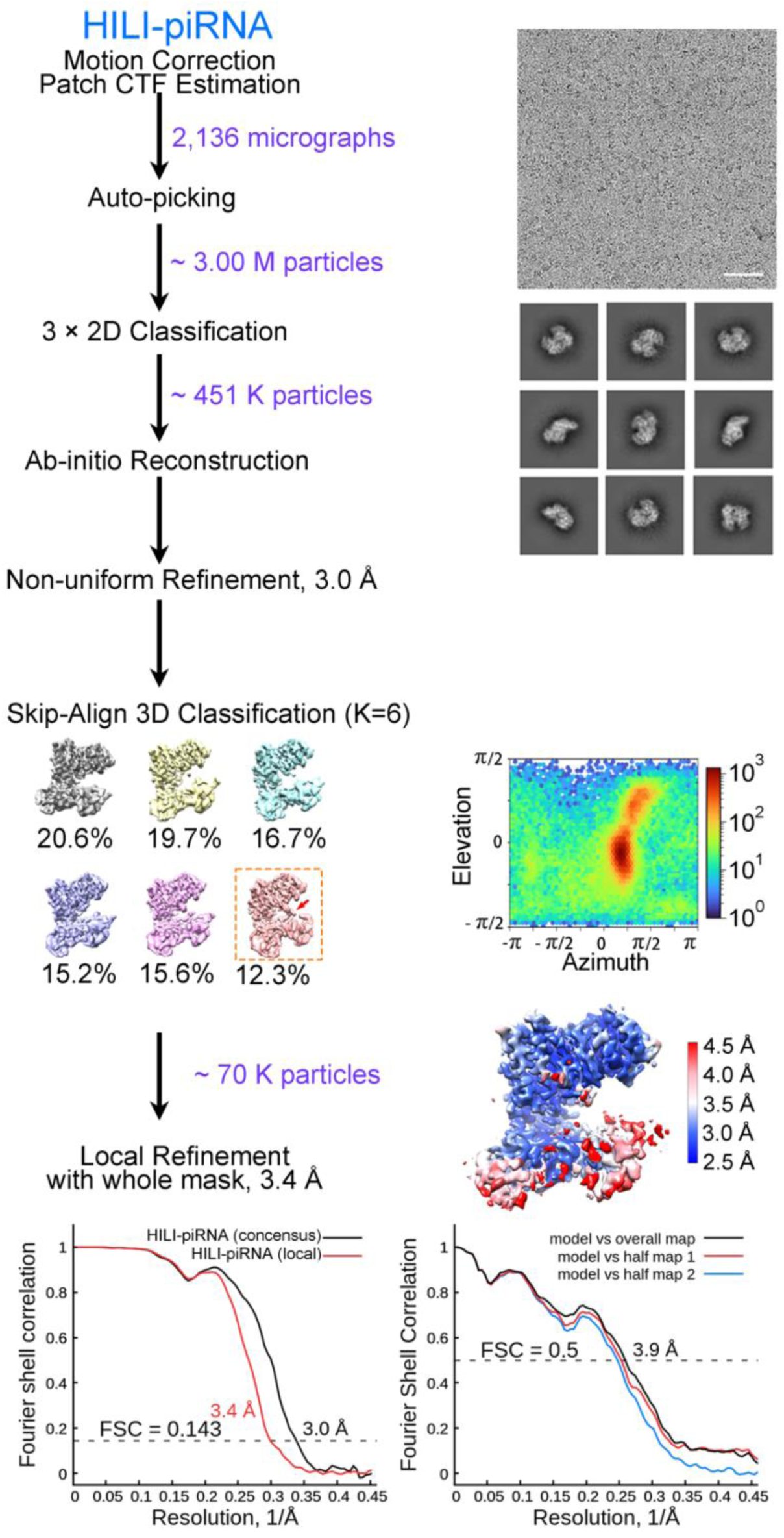
Cryo-EM analysis of HILI-piRNA. Data processing workflow for HILI-piRNA cryo-EM analysis. Representative motion-corrected dose-weighted micrograph (out of 2,136 micrographs) with a scale bar of 50 nm, 2D class averages with a box size of 209 Å, 3D classification, 3D refinement reconstruction with local resolution, angular distribution, gold-standard FSC curves of the final map (resolution cut-off at FSC = 0.143), and model vs. map curves of the final map against the final model (resolution cut-off at FSC = 0.5). 3D classes selected for subsequent analysis were indicated with dotted boxes in different colors.

**Extended Data Fig. 7.**
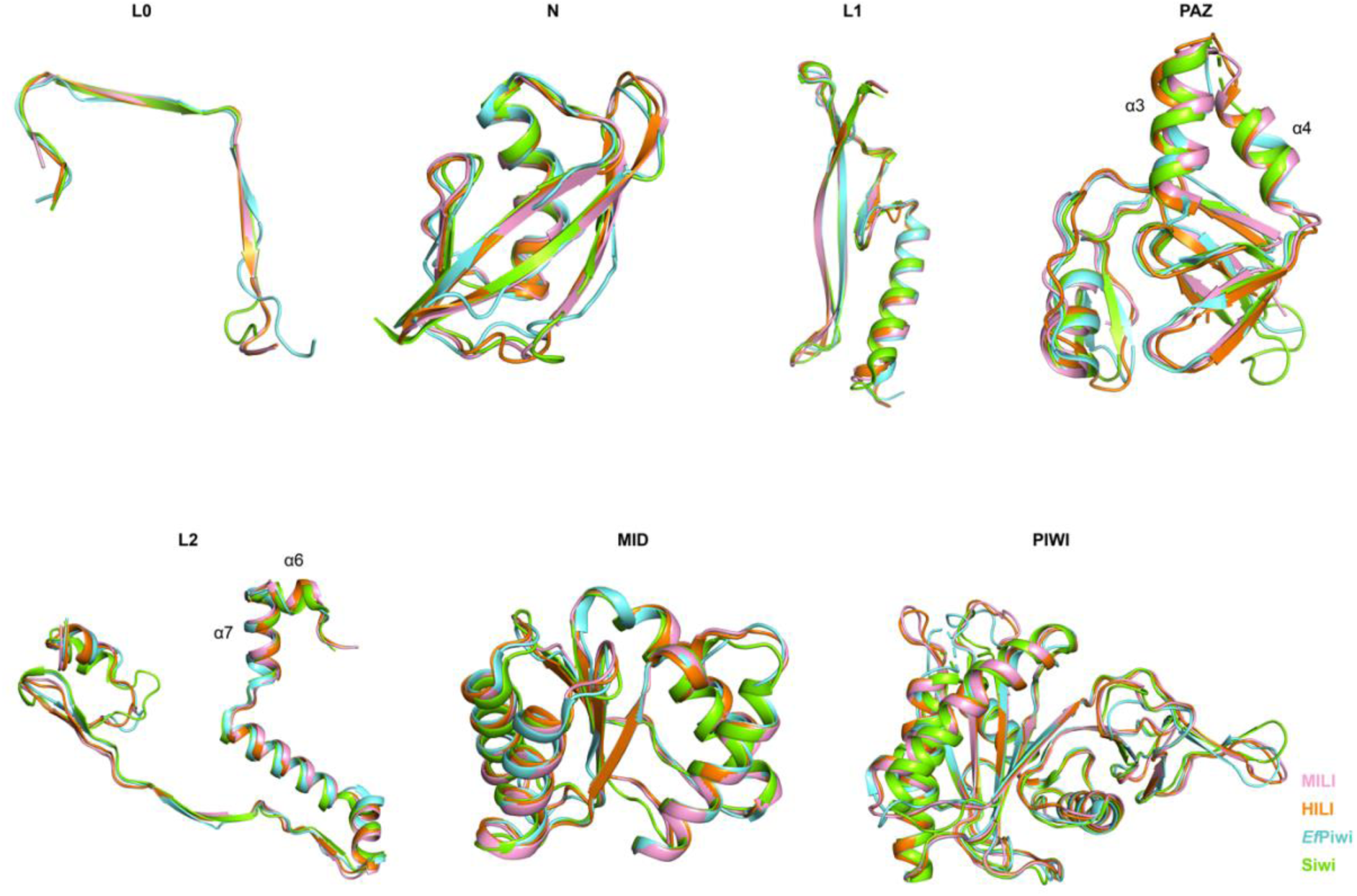
Structural comparison of individual domains of MILI, HILI, *Ef*Piwi and Siwi. Superposition of the individual domains of MILI (pink), HILI (orange), *Ef*Piwi (PDB: 7KX7) (aquamarine), and Siwi (PDB: 5GUH) (chartreuse).

**Extended Data Fig. 8.**
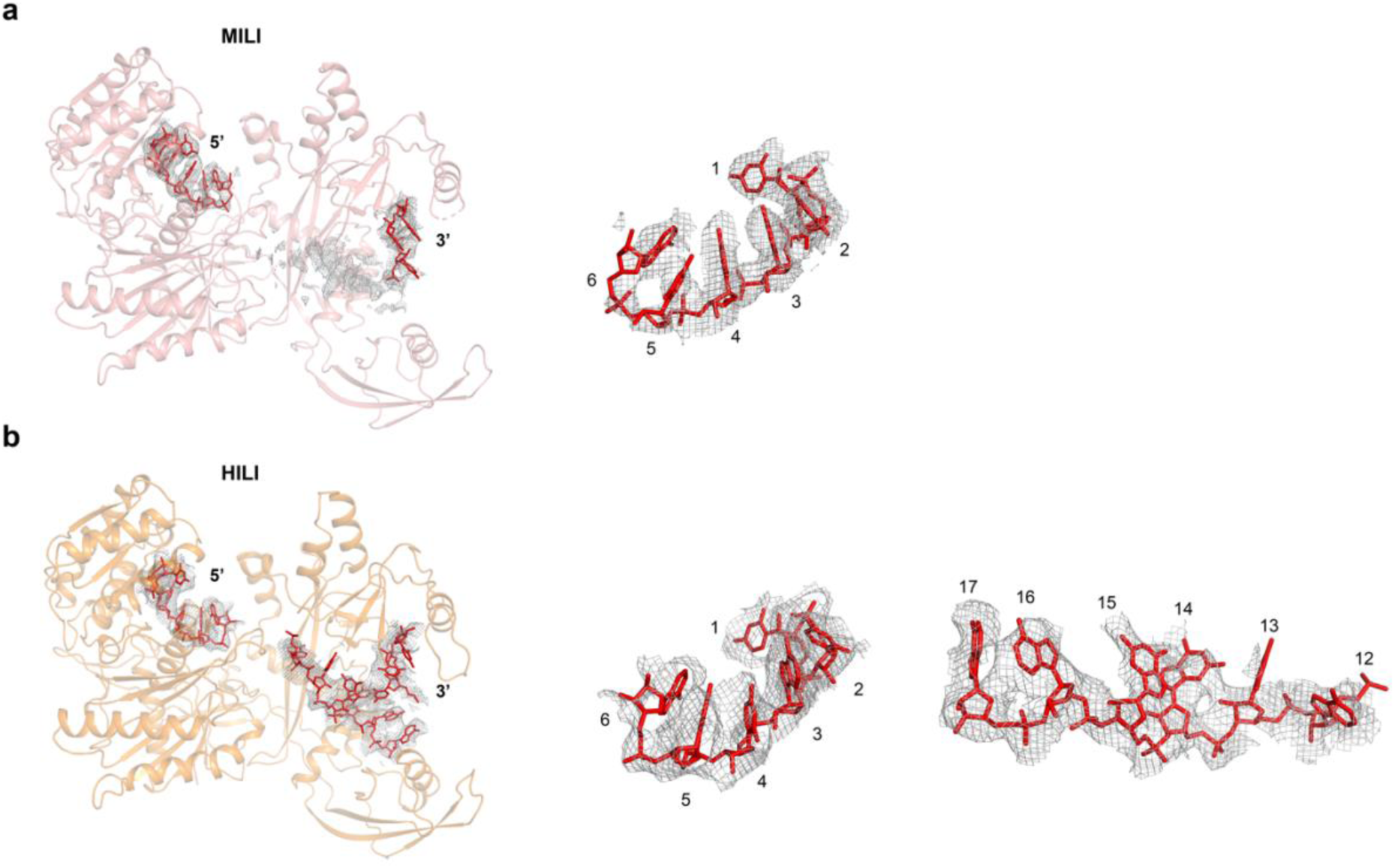
Electron density map. **a**, The density map (grey mesh) for the entire bound piRNA in MILI-piRISC is shown in the left panel. A close-up view of the EM density map (grey mesh) for the piRNA 5′ segment is shown in the right panel. Modeled guide RNAs are shown as red sticks. **b,** The density map (grey mesh) for the entire bound piRNA in HILI-piRISC is shown in the left panel. Two close-up views of the EM density map (grey mesh) for the piRNA 5′ segment and 3′ half are shown in the middle and right panels. Modeled guide RNAs are shown as red sticks.

**Extended Data Fig. 9.**
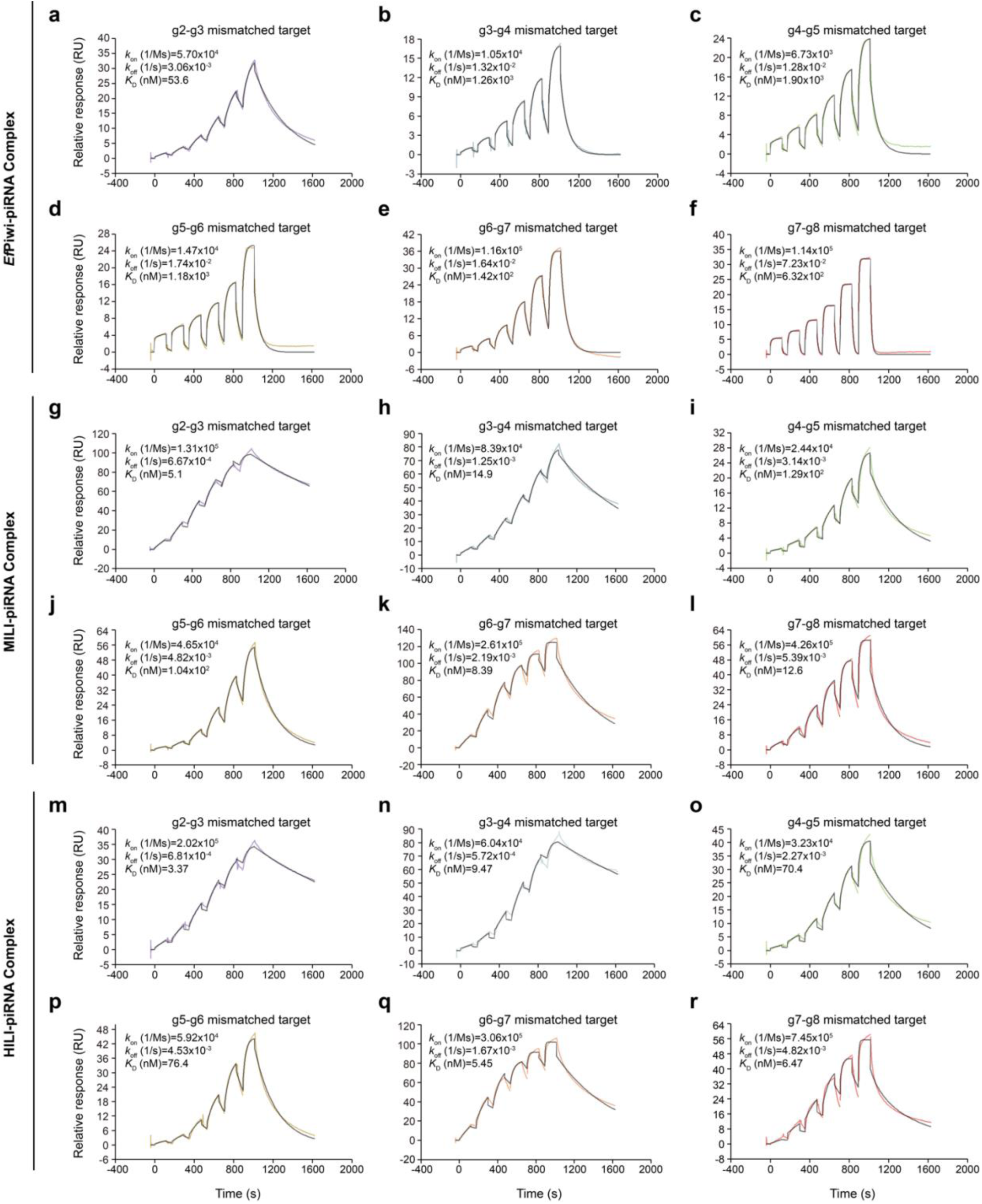
Kinetics of Piwi-piRNA complexes binding target RNAs (14 nt) with two consecutive mismatches in the seed region. **a-f**, Representative SPR traces of *Ef*Piwi-piRNA complexes binding target RNAs (14 nt) with two consecutive mismatches in the seed region, as indicated. **g-l,** Representative SPR traces of MILI-piRNA complexes binding target RNAs (14 nt) with two consecutive mismatches in the seed region, as indicated. **m-r,** Representative SPR traces of HILI-piRNA complexes binding target RNAs (14 nt) with two consecutive mismatches in the seed region, as indicated. Data shown in colors and fit the binding model shown in the grey line.

**Extended Data Fig. 10.**
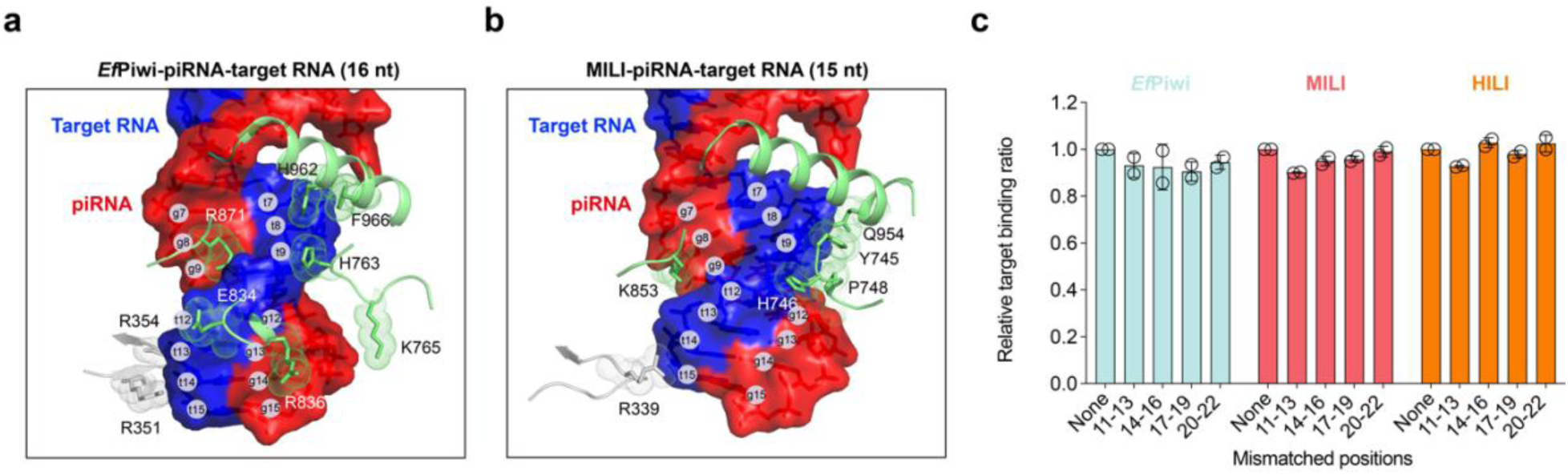
structural basis for piRNA targeting in the non-seed region. **a,b**, Non-seed interactions between piRNA-target RNA duplex and *Ef*Piwi protein (**a**) and MILI protein (**b**). **c,** Relative target-binding ratios for MILI, HILI, and *Ef*Piwi. The relative ratio here is mismatched target binding % over fully matched target binding %. Data are expressed as mean ± s.d.

**Extended Data Fig. 11.**
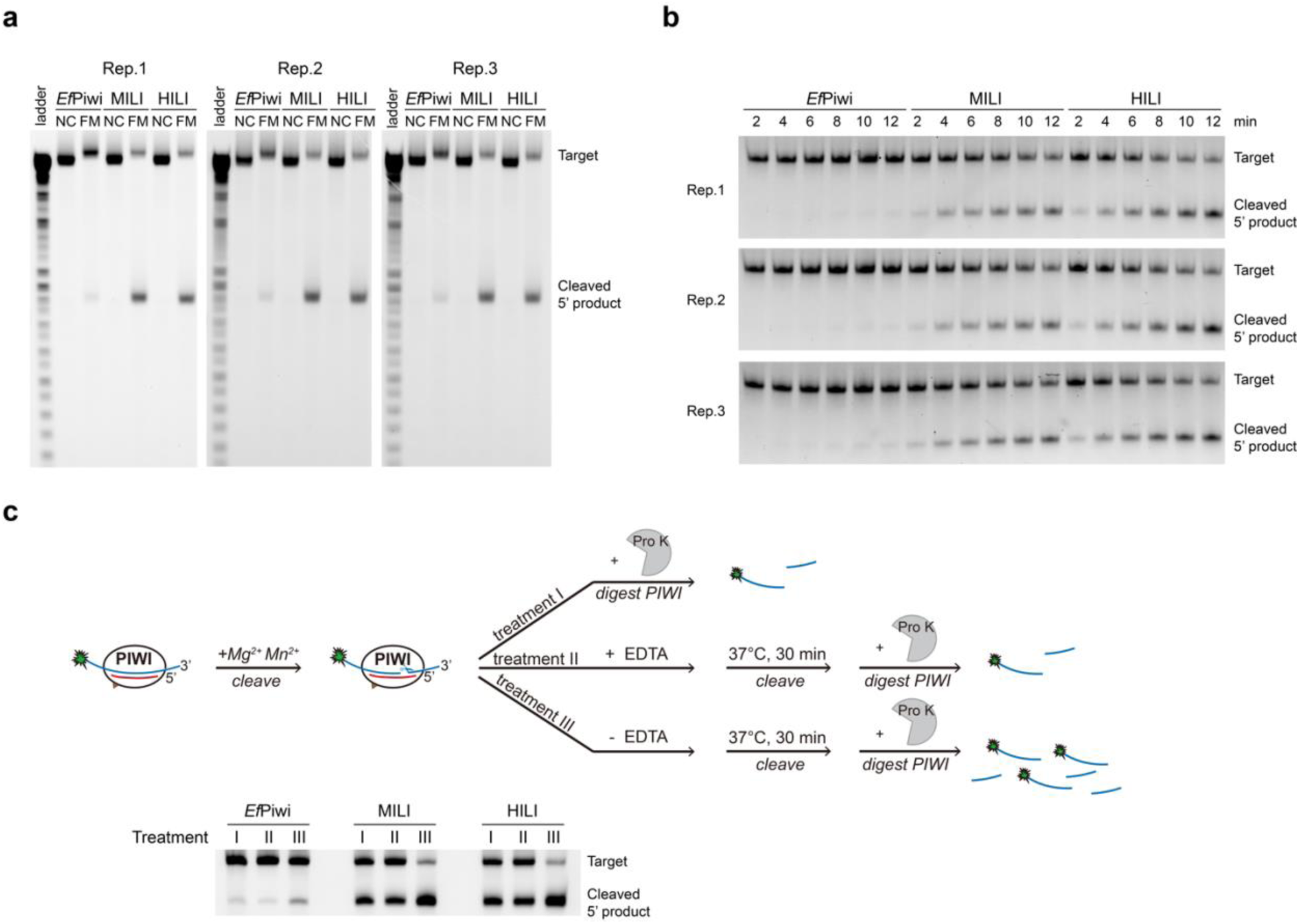
Target cleavage by *Ef*Piwi, MILI and HILI. **a**, Replicate *in vitro* cleavage reactions with equimolar amounts of substrate (target RNA) and enzyme (Piwi-piRNA complexes). NC, non-complementary target as negative control. FM, fully matched target. Reaction products were analyzed by denaturing urea-PAGE, alongside 1-nt RNA standards (left). **b,** Cleavage analysis of Piwi-piRNA complexes. Purified *Ef*Piwi-piRNA, MILI-piRNA, and HILI-piRNA (100 nM, final) were incubated with 5′-FAM-labeled target RNAs (10 nM, final) at 37°C for the indicated times. Reaction products were resolved by denaturing urea-PAGE. **c,** Effect of EDTA on chelating metal ions to stop cleavage reactions. Piwi-piRNA complexes were incubated with 5′-FAM-labeled target RNA in the presence of divalent cations for 5 mins and then were subject to three treatments. Treatment I: Protease K (Pro K) was directly added to stop the cleavage reactions. Treatment II: EDTA (5 mM, final concentration) was added to the reaction and incubated for 30 mins at 37°C before adding Pro K. Treatment III: Buffer alone was added to the reaction and incubated for 30 mins at 37°C, before adding Pro K. RNAs were analyzed by denaturing urea-PAGE.

**Extended Data Fig. 12.**
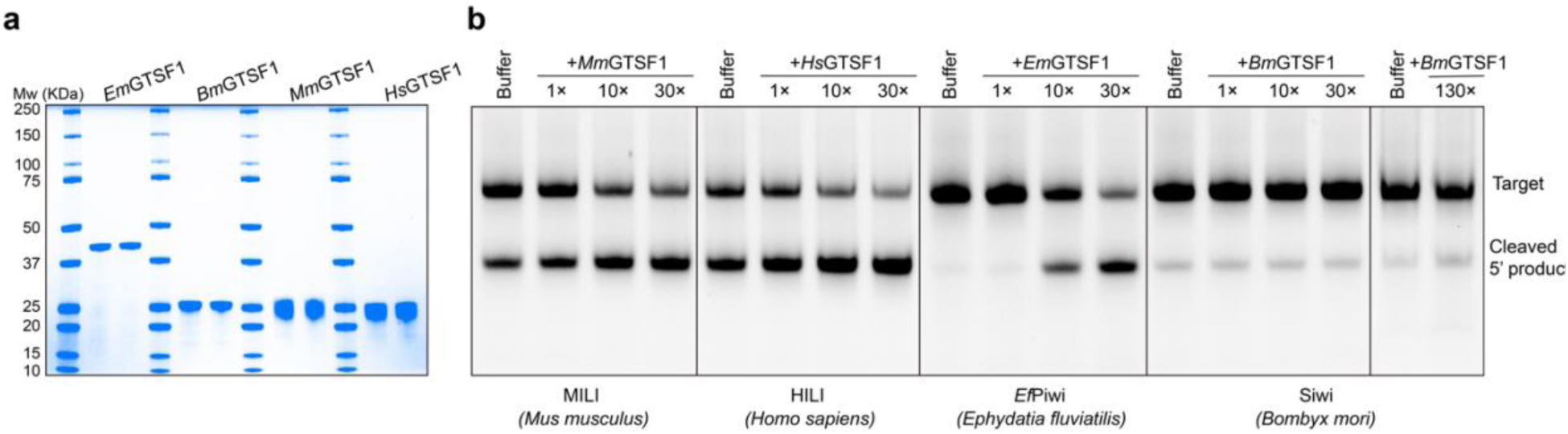
Effect of GTSF1 on slicer activity of PIWI proteins. **a**, Coomassie blue-stained SDS-PAGE of purified recombinant Flag-tagged GTSF1 proteins from different species. **b,** piRISC target cleavage assays in the presence of increasing amounts of GTSF1. *Em*, *Ephydatia muelleri*; *Bm, Bombyx mori*; *Mm*, *Mus musculus*; *Hs*, *Homo sapiens*. Note: The *Ephydatia fluviatilis* genome sequence is unavailable, so *Ef*GFST1 remains unidentified. GTSF1 from the related sponge *Ephydatia muelleri* was therefore incubated with *Ef*Piwi.

**Extended Data Fig. 13.**
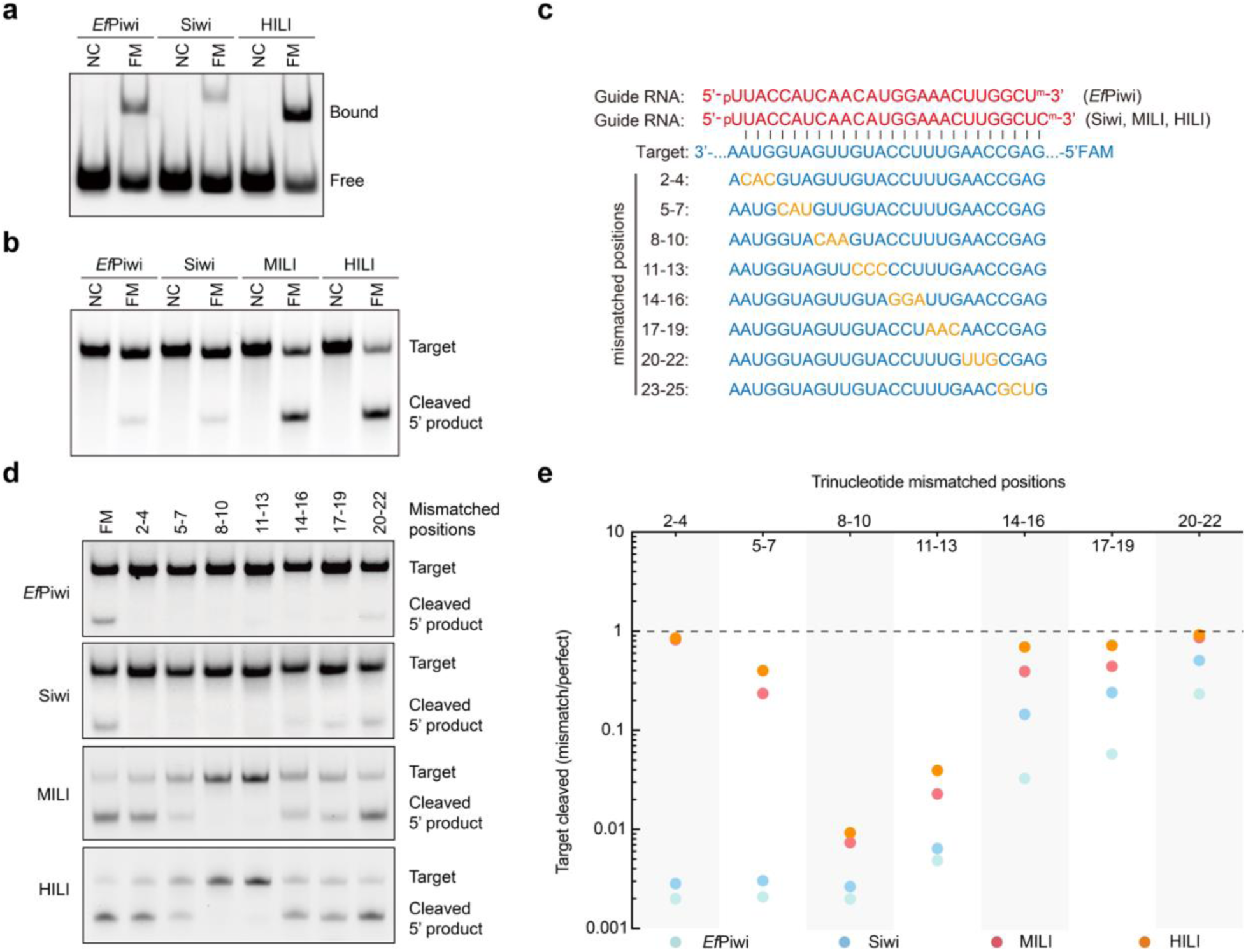
Biochemical determination of target binding, target cleavage and mismatch tolerance of Siwi. **a**, Representative native PAGE analysis of protein-guide RNA complexes binding to fully matched (FM) target RNA. NC, non-complementary (NC) target negative control. **b,** Representative *in vitro* cleavage reactions in the presence of equimolar substrate (target RNA) and enzyme (Piwi-piRNA complexes). **c,** Guide-target-pairing schematic for targets with 3-nt mismatch regions to guide RNA used in (**d, e**). **d,** Representative urea-PAGE showing the ability of *Ef*Piwi, Siwi, MILI, and HILI to cleave targets with three consecutive mismatches between g2 and g22. **e,** Quantification of the *in vitro* cleavage assays (right) showing changes in cleavage activity for three consecutive mismatches between g2 and g22.

**Extended Data Fig. 14.**
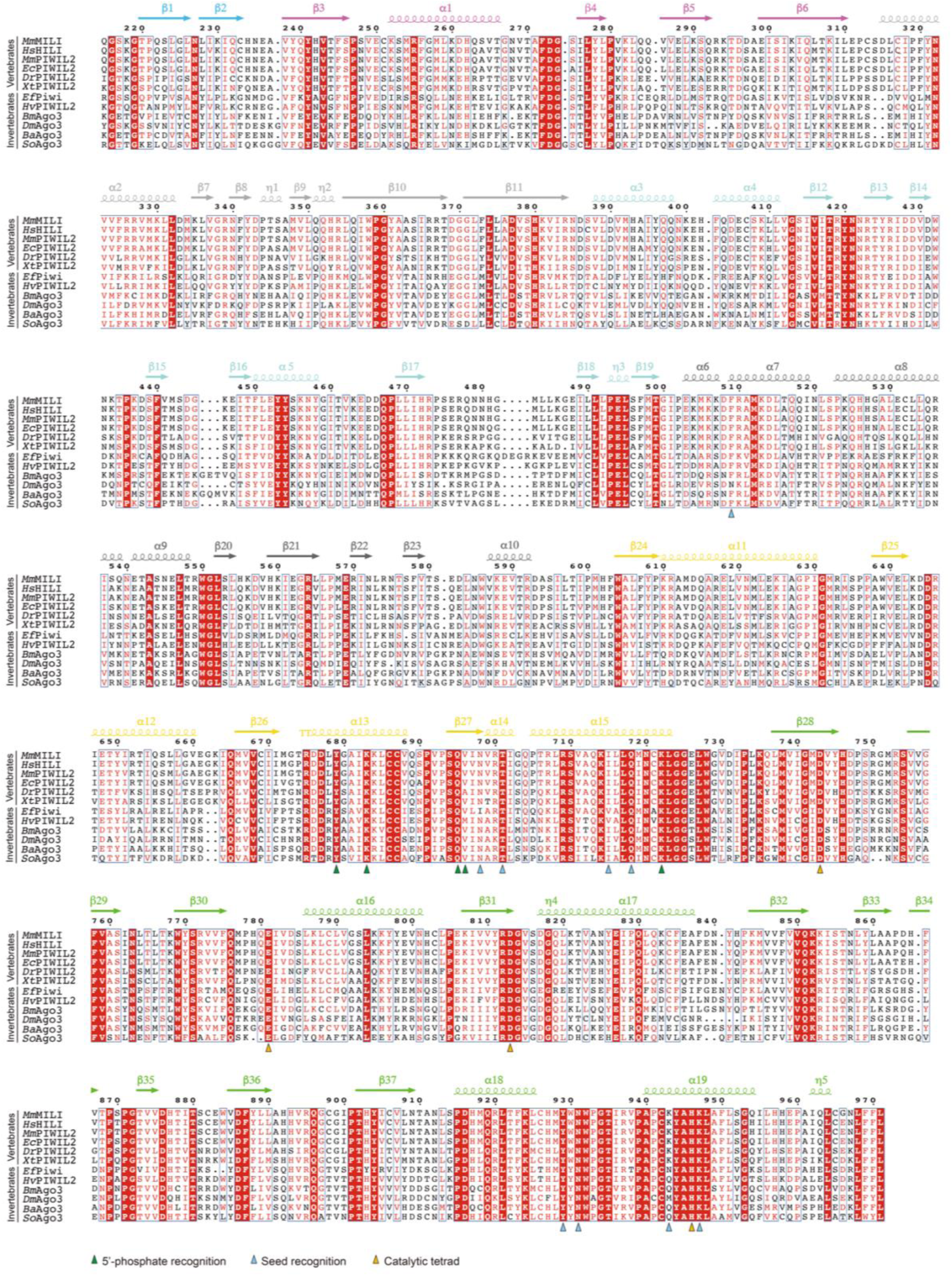
Sequence alignment of vertebrate Ago3-like PIWI proteins. The secondary structure of *Mm*MILI is indicated above the sequences, color coded by domains as indicated in Fig. 2. Key residues are marked by triangles. Species abbreviations: *Mm*, *Mus musculus* (mouse); *Hs*, *Homo sapiens* (human); *Mm, Macaca mulatta* (macaque); *Ec, Equus caballus* (horse); *Dr, Danio rerio* (zebrafish); *Xt, Xenopus tropicalis* (frog); *Ef, Ephydatia fluviatilis* (sponge); *Hv, Hydra vulgaris* (hydra); *Bm, Bombyx mori* (silkworm); *Dm, Drosophila melanogaster* (fruit fly); *Ba, Bicyclus anynana* (wood nymph); *So, Sitophilus oryzae* (rice weevil).

**Extended Data Fig. 15.**
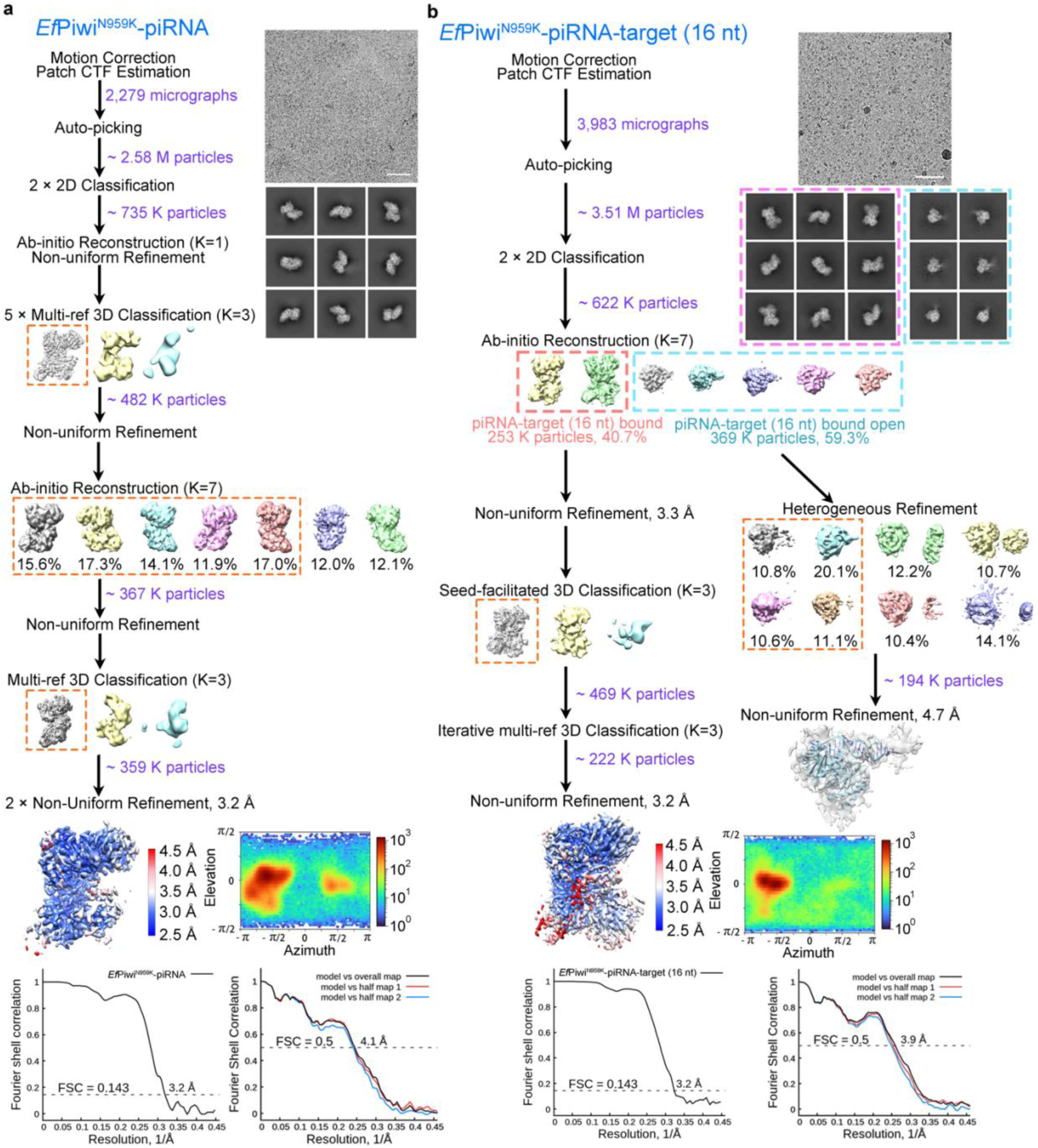
Cryo-EM analysis of *Ef*Piwi^N959K^-piRNA, *Ef*Piwi^N959K^-piRNA-target (16 nt). **a,b,** Data processing workflow of cryo-EM analysis of *Ef*Piwi^N959K^-piRNA (**a**) and *Ef*Piwi^N959K^-piRNA-target (16 nt) (**b**). Representative motion-corrected dose-weighted micrograph (out of 2,279, 3,983 micrographs) with a scale bar of 50 nm, 2D class averages with a box size of 209 Å, 3D classification, 3D refinement reconstruction with local resolution, angular distribution, gold-standard FSC curves of the final map (resolution cut-off at FSC = 0.143), and model vs. map curves of the final map against the final model (resolution cut-off at FSC = 0.5). 3D classes selected for subsequent analysis were indicated with dotted boxes in different colors.

**Extended Data Fig. 16.**
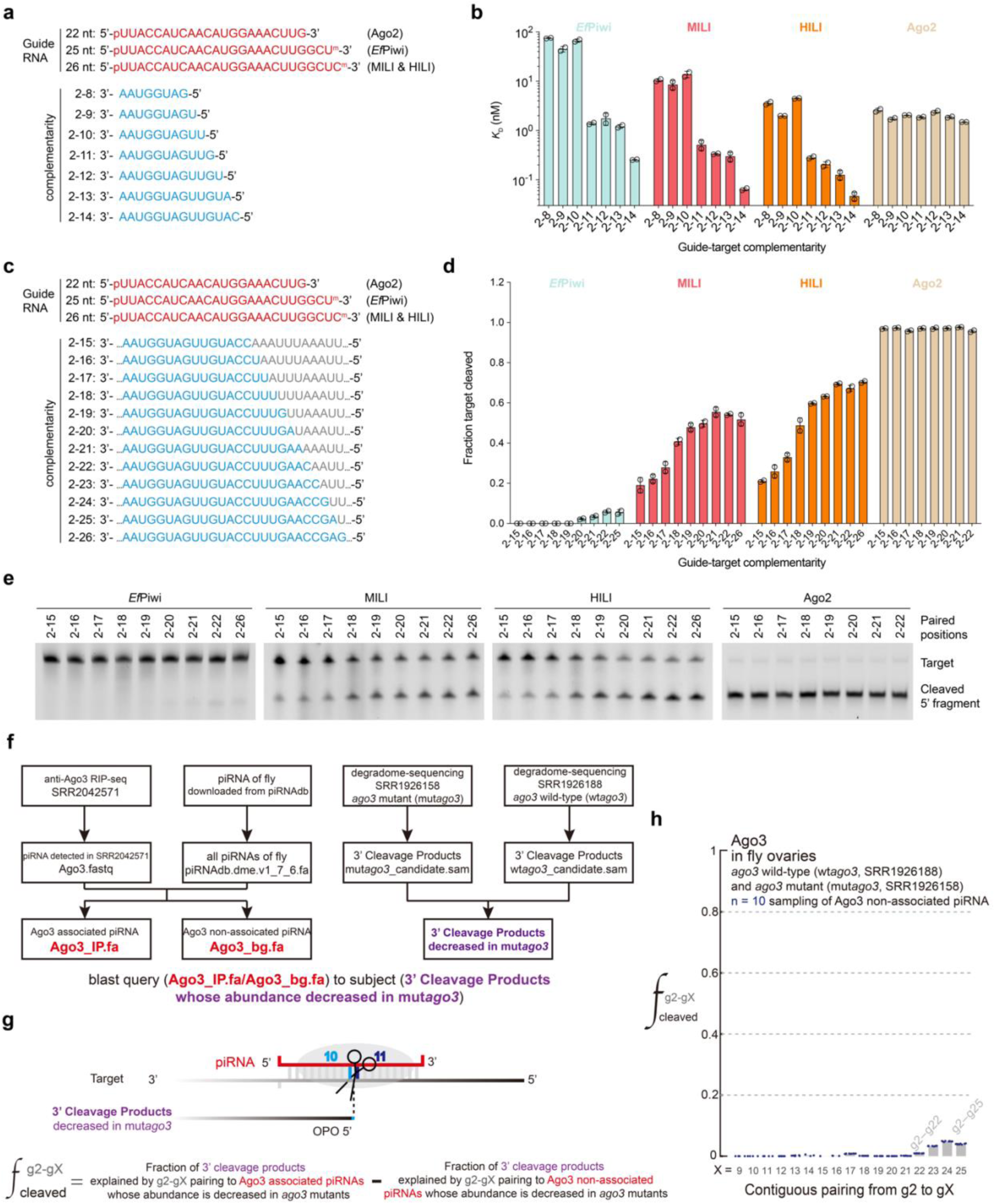
Target binding and cleavage data by *Ef*Piwi, MILI, HILI and Ago2 *in vitro* and analysis of *Drosophila* Ago3-catalyzed splicing of transcripts *in vivo*. **a**, Guide-target pairing schematic for targets with increasing complementarity for target binding analysis in (**b**). **b,** Equilibrium dissociation constants (*K*_D_) of Argonaute-guide complexes binding target RNAs with increasing complementarity. **c,** Guide-target pairing schematic for targets with increasing complementarity for cleavage analysis in (**d, e**). **d,** Quantification of reaction products shown as fraction of target cleaved. **e,** Representative urea-PAGE showing *Ef*Piwi-, MILI-, HILI-, and Ago2-mediated cleavage of targets with increasing guide complementarity. In (**b**) and (**d**), data are represented as mean ± s.d. **f,** Schema of data preparation and preprocessing to identify Ago3-mediated cleavage products in *Drosophila* ovaries. Ago3-bound piRNAs (Ago3_IP.fa) and non-associated piRNA (Ago3_bg.fa) were identified and compared to potential 3′ cleavage products whose abundance decreased in mutant *ago3* degradome sequencing libraries (mut*ago3*; SRR1926158) compared to wild-type *ago3* degradome sequencing libraries (wt*ago3*; SRR1926188). See Methods for details. **g,** piRNA queries (Ago3_IP.fa/Ago3_bg.fa) were aligned to candidate targets whose cleaved 5′ ends are complementary to g2–g10 of a piRNA. Targets with extended pairing were identified from g2–g25. For each pairing arrangement g2–gX, the fraction of cleaved targets was calculated as the fraction of cleaved targets explained by Ago3-associated piRNAs minus the fraction explained by Ago3-non-associated piRNAs. **h,** Plot of the fraction of cleaved targets in fly ovaries for contiguous pairing from nucleotide g2–gX, as indicated. Ago3-non-associated piRNA were sampled 10 times.

**Extended Data Table 1.**
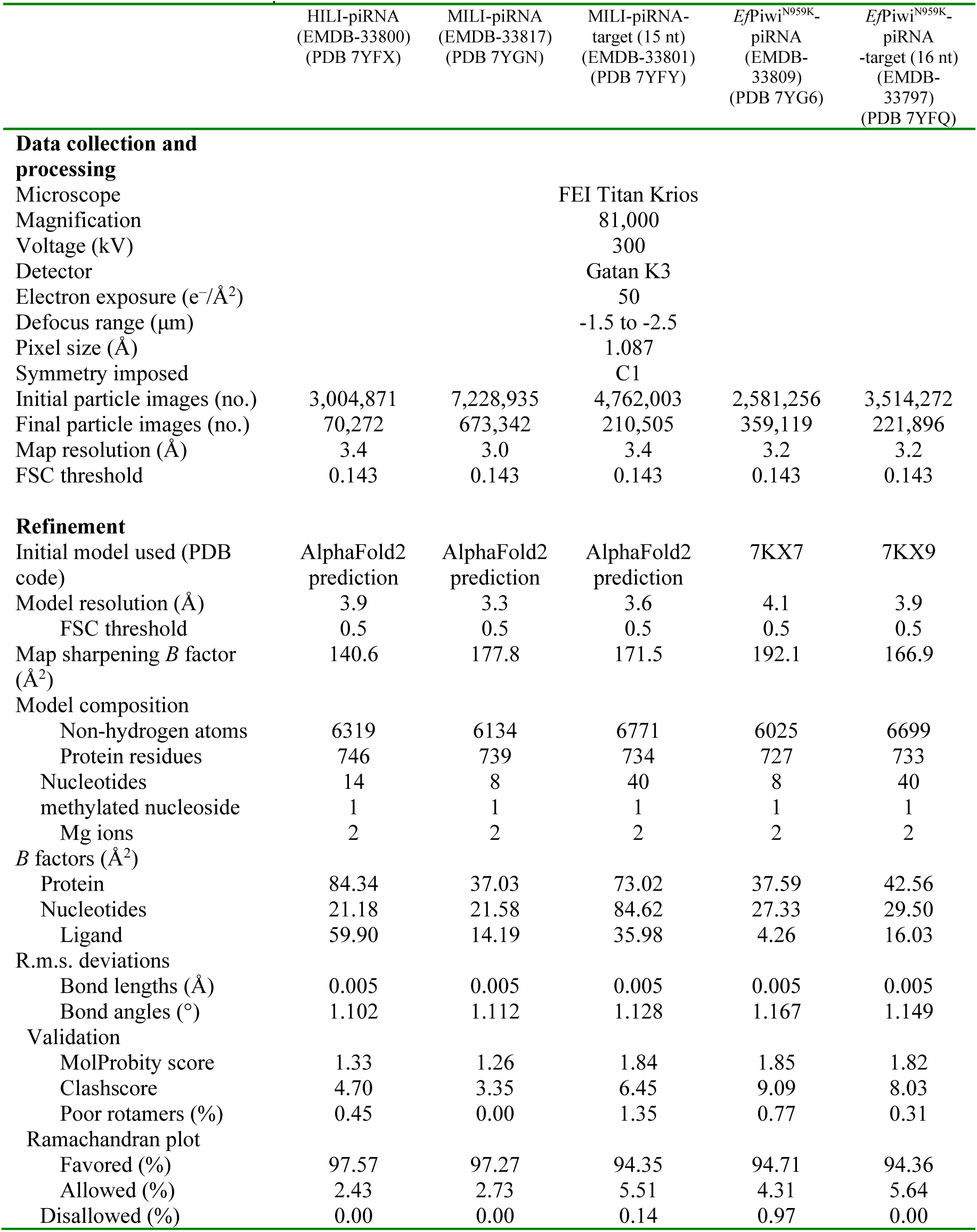
Statistics for data collection and structural refinement.

**Extended Data Table 2.**
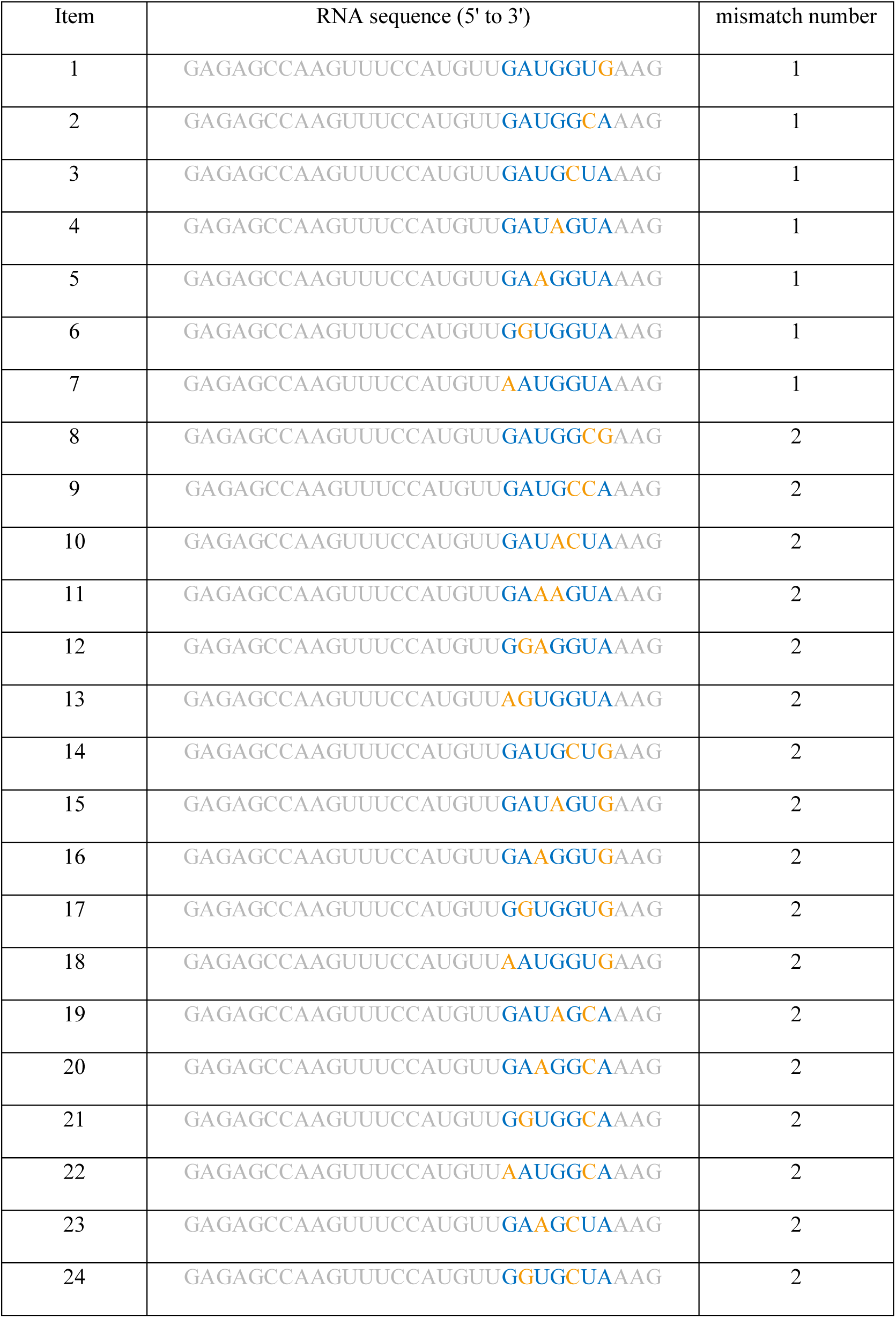
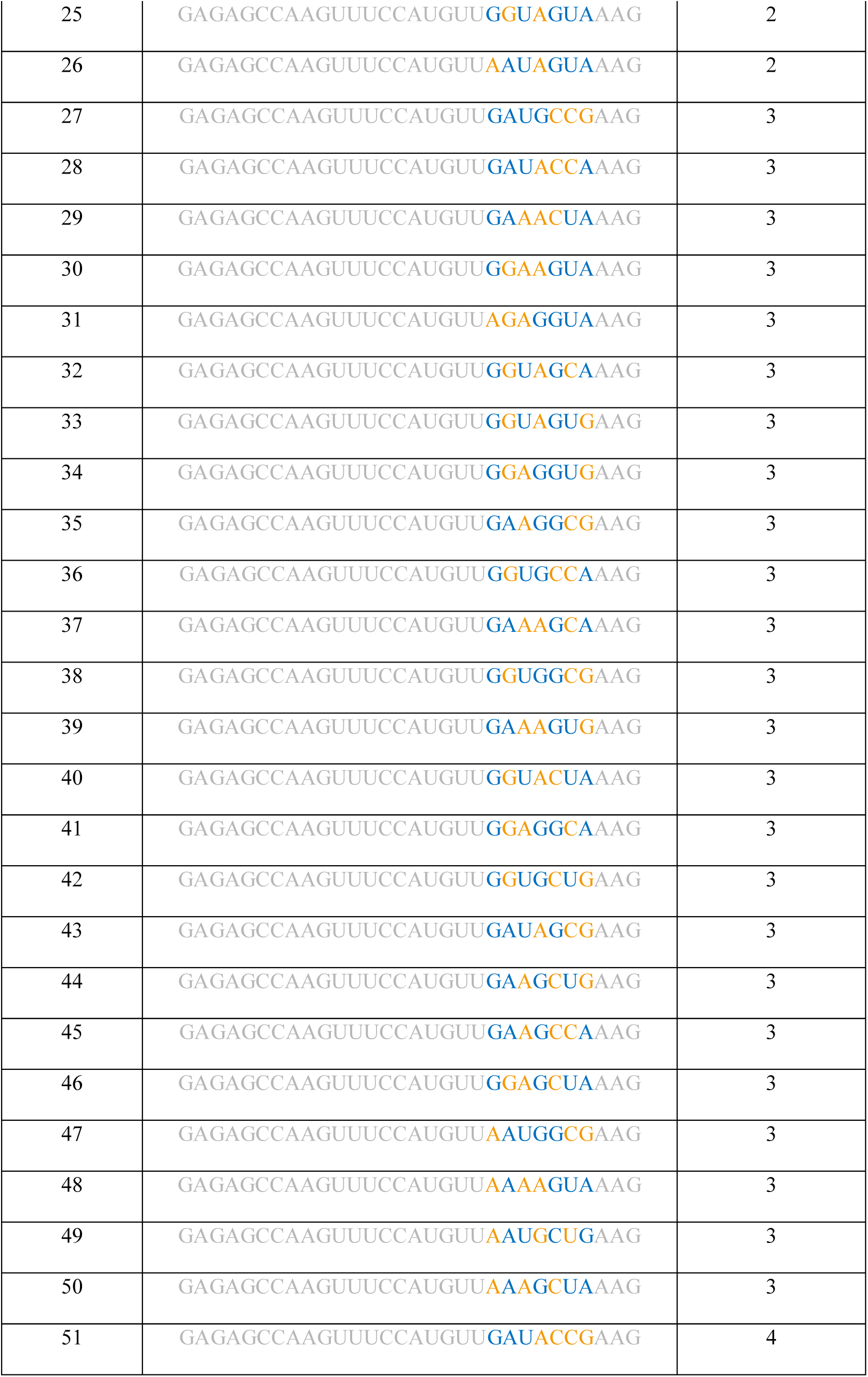
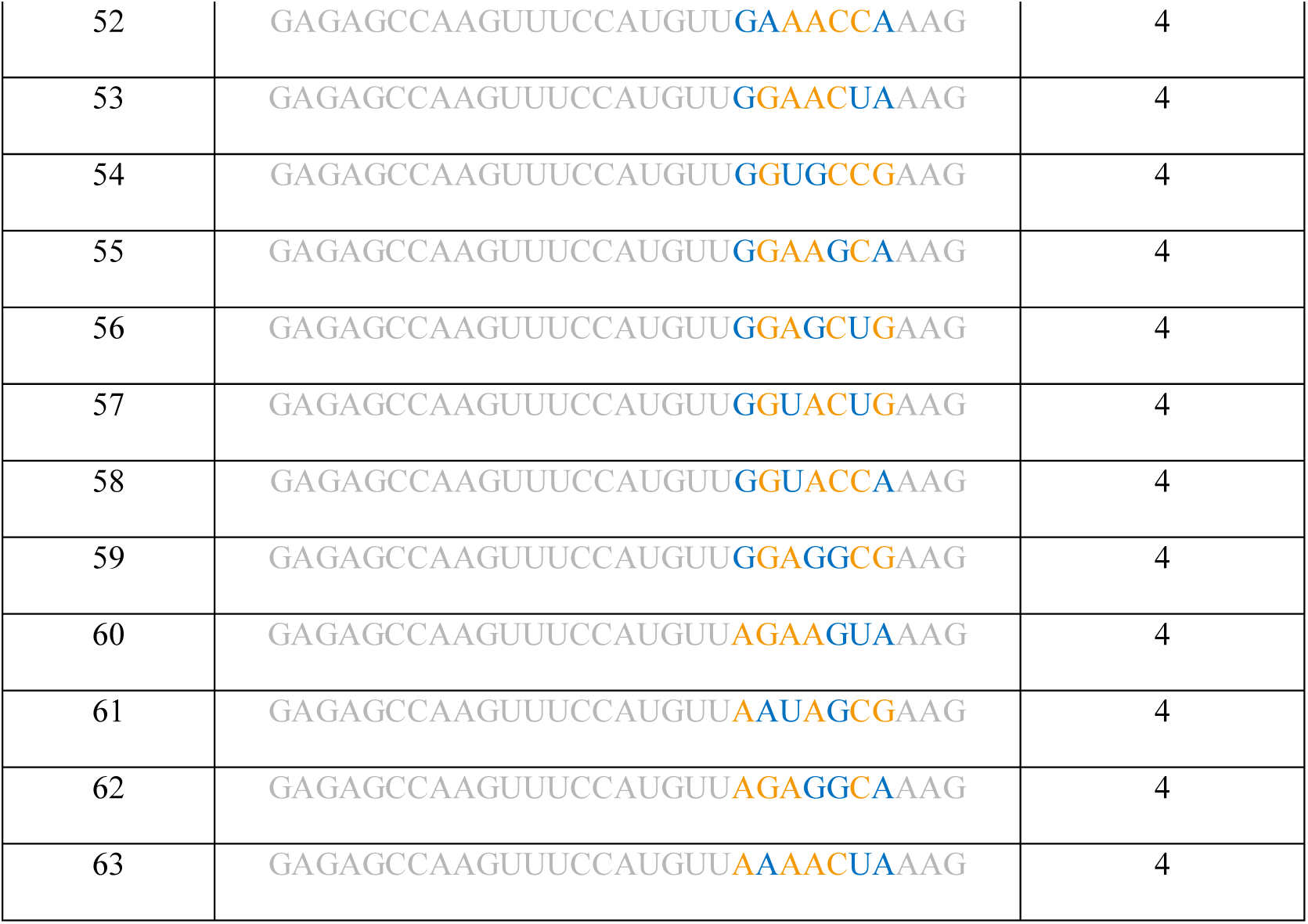
Sequences of the 63 target RNAs, with 1-4 mismatches opposite piRNA nucleotides g2-g8.

